# Inhibition is the hallmark of CA3 intracellular dynamics around awake ripples

**DOI:** 10.1101/2021.04.20.440699

**Authors:** Koichiro Kajikawa, Brad K. Hulse, Athanassios G. Siapas, Evgueniy V. Lubenov

## Abstract

Hippocampal ripples are transient population bursts that structure cortico-hippocampal communication and play a central role in memory processing. However, the mechanisms controlling ripple initiation in behaving animals remain poorly understood. Here we combine multisite extracellular and whole cell recordings in awake mice to contrast the brain state and ripple modulation of subthreshold dynamics across hippocampal subfields. We find that entorhinal input to DG exhibits UP and DOWN dynamics with ripples occurring exclusively in UP states. While elevated cortical input in UP states generates depolarization in DG and CA1, it produces persistent hyperpolarization in CA3 neurons. Furthermore, growing inhibition is evident in CA3 throughout the course of the ripple buildup, while DG and CA1 neurons exhibit depolarization transients 100 ms before and during ripples. These observations highlight the importance of CA3 inhibition for ripple generation, while pre-ripple responses indicate a long and orchestrated ripple initiation process in the awake state.

## INTRODUCTION

Bidirectional interactions between the hippocampus and neocortical areas are believed to play a key role in memory consolidation (Squire, 1992). Hippocampal ripples are deemed essential for this process because the associated population activity reflects prior experience (Foster, 2017; Kudrimoti et al., 1999; Lee and Wilson, 2002; Wilson and McNaughton, 1994) and ripple disruption results in memory deficits (Ego-Stengel and Wilson, 2010; Girardeau et al., 2009; Jadhav et al., 2012). Ripples provide synchronous volleys that drive cortical targets and co-occur with distinct cortical network patterns (Battaglia et al., 2004; Jiang et al., 2019; Ji and Wilson, 2007; Logothetis et al., 2012; Mölle et al., 2006; Shein-Idelson et al., 2016; Siapas and Wilson, 1998; Wierzynski et al., 2009). In particular, ripples normally occur during slow-wave sleep (SWS) and quiet wakefulness when hippocampal local field potentials (LFPs) display large-amplitude irregular activity (LIA) (Buzsáki, 1986; Jarosiewicz and Skaggs, 2004; Kay et al., 2016; O’Keefe, 1976; Vanderwolf, 1969), whereas neocortical dynamics are marked by the presence of UP and DOWN states (UDS), alternating periods of elevated and depressed network activity that can also be observed under anesthesia (Cowan and Wilson, 1994; Steriade et al., 1993b, 1993c). Neocortical and hippocampal dynamics can be coordinated via the entorhinal cortex (EC), the main gateway between neocortical areas and the hippocampus, which provides direct input to the dentate gyrus (DG), and areas CA3 and CA1 (Amaral and Witter, 1989). Experiments in sleeping and anesthetized animals show that the EC also exhibits UDS that modulate activity across hippocampal subfields (Hahn et al., 2012; Isomura et al., 2006). However, the influence of cortical UDS on hippocampal dynamics and ripple generation in wakefulness are not well understood.

Ripples are believed to be the product of excitatory buildup in the recurrent CA3 network, culminating in a population burst that drives CA1 spiking organized by the transient ripple oscillation (Buzsáki, 1986; Miles and Wong, 1983; Stark et al., 2014; Traub and Miles, 1991). This suggests that CA3 neurons should get progressively depolarized and come closer to firing threshold through the course of the ripple buildup. Separate from the buildup itself, the processes controlling ripple initiation and termination are not fully understood. *In vitro* experiments indicate that ripples are initiated once stochastic fluctuations in the population firing rate of CA3 pyramidal cells exceed a threshold level (de la Prida et al., 2006; Schlingloff et al., 2014), implying that an increase in the firing of CA3 neurons should result in a corresponding increase in the rate of ripple occurrence. Recent *in vitro* studies have also emphasized the importance of inhibitory neurons in ripple initiation (Bazelot et al., 2016; Ellender et al., 2010; Schlingloff et al., 2014), while other studies suggested that area CA2 and a special class of CA3 cells play a key role in ripple initiation (Hunt et al., 2018; Oliva et al., 2016). Furthermore, there is growing evidence that the functional role of ripples as well as the mechanism of their initiation may differ across the awake and sleep states (Middleton and McHugh, 2020; Oliva et al., 2016; Roumis and Frank, 2015; Tang and Jadhav, 2019). Whole cell recordings in awake animals have opened a window to understanding the interplay between collective network activity and membrane potential (V_m_) dynamics of hippocampal neurons which can reveal the nature and timing of synaptic inputs and subthreshold changes that are invisible to extracellular recordings. While recent efforts have examined the V_m_ of CA1 neurons around ripples (English et al., 2014; Hulse et al., 2016) and the V_m_ modulation by brain state across hippocampal subfields (Hulse et al., 2017; Malezieux et al., 2020), the subthreshold dynamics of CA3 pyramidal cells around ripples and the impact of cortical inputs on V_m_ behavior in CA3 remain unknown.

Here we combine multisite extracellular and whole cell recordings in awake mice to characterize and contrast the membrane potential dynamics of principal neurons in CA3 with that in the dentate gyrus (DG) and CA1. Our results reveal that the membrane potential of CA3 neurons is modulated by brain state and ripples in a way that is largely opposite to the modulation of dentate granule cells and CA1 neurons. This divergent modulation across hippocampal subfields is surprising given that they are interconnected and receive similar inputs from the entorhinal cortex, and offers insights into the processes of ripple initiation, build up, and termination.

## RESULTS

We combined whole-cell recordings of principal neurons across DG, CA3, and CA1 with multisite extracellular local field potential (LFP) recordings spanning the radial extent of dorsal CA1 and DG in awake head-fixed mice that were free to run on a spherical treadmill (Figure 1). Below we show that entorhinal inputs to DG exhibit UP and DOWN dynamics during quiet wakefulness, with ripples occurring exclusively in the UP state. Analysis of how these brain states influence membrane potential dynamics, reveals that CA3 neurons hyperpolarize in the UP state when ripples occur, in contrast to neurons in DG and CA1. We then focus on characterizing how brain state is reflected in slow V_m_ trends around ripples. Finally, we analyze ripple triggered modulation of fast V_m_ fluctuations which reveals a prevalence of inhibition in CA3 which grows through the ripple buildup. In contrast DG and CA1 neurons exhibit transient depolarization not only after ripple onset but also 100 ms earlier.

**Figure 1.**
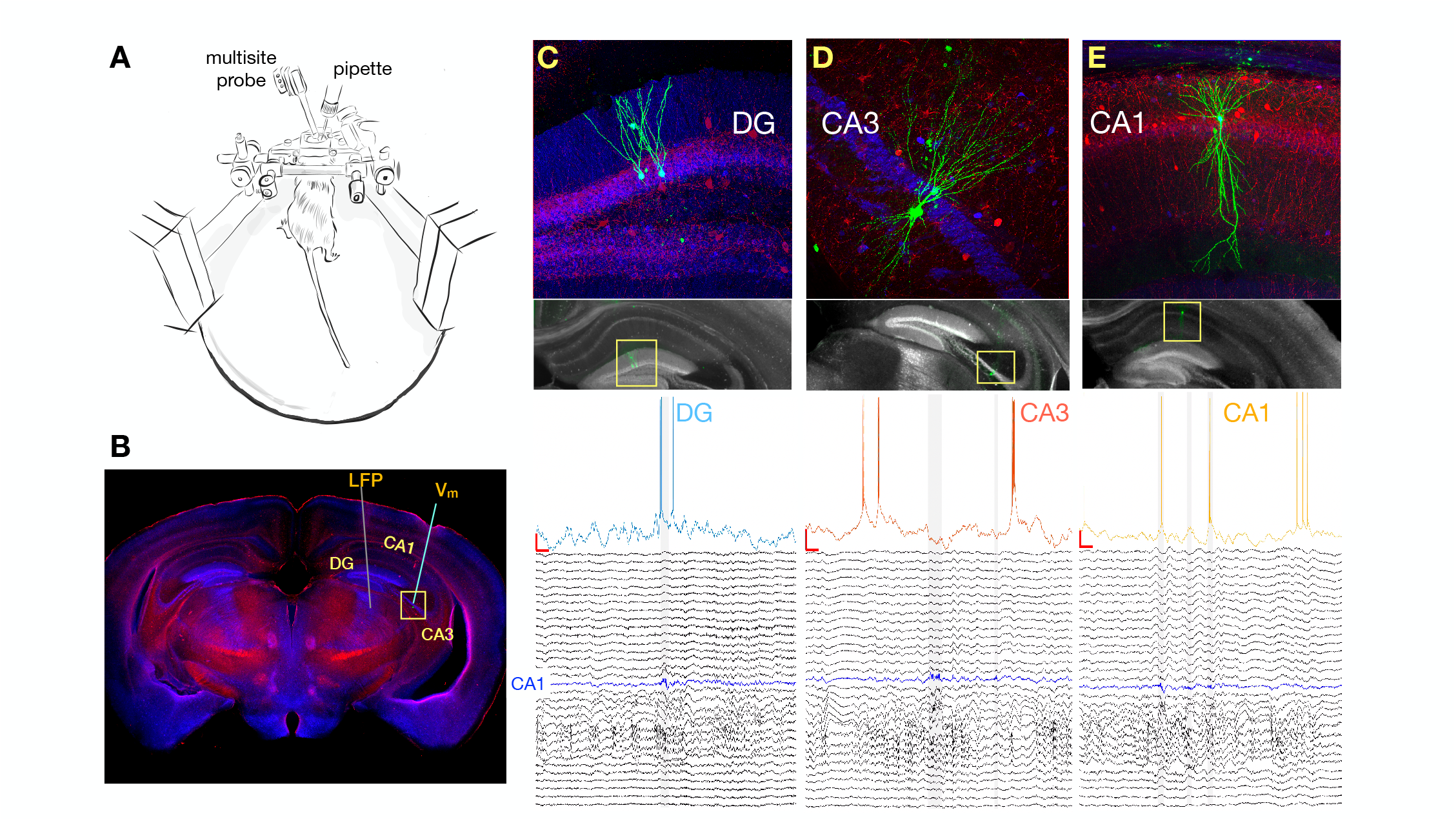
Simultaneous Multisite Extracellular and Whole-Cell Recordings Across Hippocampal Subfields. **(A)** Schematic of setup for simultaneous intracellular and extracellular recordings from awake head-fixed mice free to run on a spherical treadmill. **(B)** Typical penetration paths of multisite probe for LFP recordings and micropipette (targeting CA3 in this example for the neuron shown in D) for whole cell recordings. Histological sections were stained for biocytin (green), calbindin (blue), and parvalbumin (red). **(C)** Top: Recorded DG granule cells are labeled with biocytin and their location and morphology is visualized with fluorescence microscopy. Bottom: Membrane potential (V_m_) from one of the above cells together with simultaneous LFP recordings spanning both CA1 and DG. The blue trace marks the pyramidal cell layer of CA1 where ripples are detected and marked by the gray vertical shading. Notice that the cell fires right before the onset of a ripple. **(D)** Same as C, but for a pyramidal neuron in CA3. Notice that the cell is hyperpolarized following the ripple onset. **(E)** Same as C, but for a pyramidal neuron in CA1. Notice that the cell fires inside two nearby ripples.

### Entorhinal Inputs to DG Exhibit UP and DOWN Dynamics During Quiet Wakefulness

Based on LFP dynamics, the network state of the hippocampus can be classified as large irregular activity (LIA), small irregular activity (SIA), or theta rhythmic, with LIA and SIA being the predominant states during quiet wakefulness (Buzsáki, 1986; Jarosiewicz and Skaggs, 2004; Kay et al., 2016; O’Keefe, 1976; Vanderwolf, 1969). Since this classification is typically based on a single hippocampal LFP, it does not consider the origin of the observed field fluctuations, but only their amplitude and frequency content. As a consequence, synaptic currents due to inputs from the entorhinal cortex (EC) as well as other hippocampal subfields are reflected in the local field and cannot be dissociated. In order to address this, we used multisite LFP recordings and computed the laminar current source density (CSD) (Mitzdorf, 1985; Pettersen et al., 2006) throughout CA1 and DG, which in combination with the known circuit anatomy allowed us to infer the spatiotemporal pattern of synaptic activity. An example estimate of this laminar CSD is illustrated in Figure 2, which shows several sharp wave-ripples (SWRs) associated with pronounced current sinks in stratum radiatum (sr) of CA1 due to CA3 synaptic input, occurring against a background of synaptic activity in DG. On a longer timescale the CSD clearly reveals alternating periods of high and low synaptic activity lasting several seconds (Figure 2B) that are particularly prominent within the DG. Strikingly, these alternating periods appear to coincide with slow shifts in the membrane potential of an example CA3 pyramidal neuron (Figure 2B). To quantify the level of cortical input to the hippocampus, we averaged the rectified CSD over the molecular layer of DG (Figure 2). This measure reflects the magnitude of synaptic currents due to inputs from layer II of the lateral and medial EC arriving at the outer two thirds of the dentate molecular layer (Sullivan et al., 2011), as well as associational and return currents flowing in the inner third. Confirming our observations, the DG CSD magnitude showed strong coherence with the subthreshold membrane potential of the example CA3 neuron for frequencies below 1 Hz (Figure 2-figure supplement 1B).

**Figure 2.**
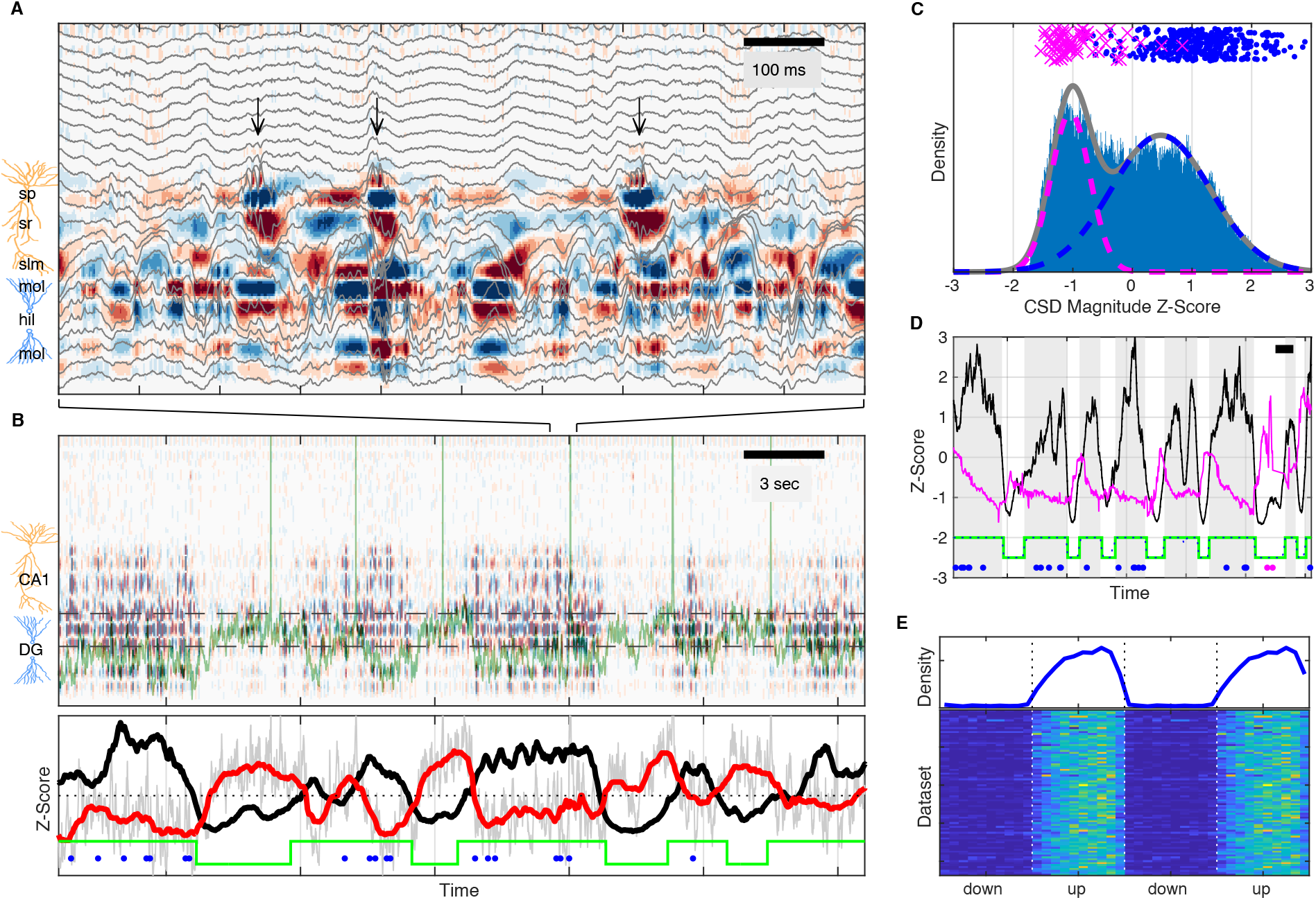
UP and DOWN States Modulate Slow V_m_ Shifts and Ripple Occurrence. **(A)** Image of current source density (CSD) derived from the LFP traces (gray). Ripples, high-frequency oscillations indicated by the black arrows, are associated with current sinks (red) in stratum radiatum (sr) below the pyramidal cell layer (sp). The bottom third of the image shows large current sources (blue) and sinks (red) within the molecular layers (mol) of DG. **(B)** Top: Image of the CSD on a longer timescale reveals alternating periods of high and low CSD activity. The two interrupted black lines mark the vertical extent of the suprapyramidal molecular layer of DG. The V_m_ of a CA3 neuron is superimposed in green. Notice that periods of low DG CSD magnitude (light colors) are associated with V_m_ depolarization. Bottom: DG CSD magnitude (black), quantified by averaging the rectified CSD over the molecular layer of DG, normalized to a z-score. Subthreshold V_m_ (gray) for the CA3 neuron and its slow component (red) plotted as z-scores. Notice that the black and red traces are anti-correlated. The green stairstep trace marks epochs of elevated DG CSD magnitude (UP states) and decreased DG CSD magnitude (DOWN states). Ripples (blue dots) occur in the UP state. **(C)** Distribution of DG CSD magnitude fitted with a two component gaussian mixture. DG CSD magnitude at ripples (blue dots) and eye blinks (magenta), show preferential association with the UP and DOWN components, respectively. **(D)** Hidden Markov model (HMM) state detection based on DG CSD magnitude (black). DG current magnitude is high in the UP state (gray stripes), while the pupil (diameter in magenta) dilates at the onset of the DOWN state and then gradually constricts in the course of the UP state. Ripples and eye blinks are marked by blue and magenta dots, respectively. Horizontal scale bar is 3 seconds long. **(E)** (Top) Population average probability density of ripple occurrence as a function of UDS phase. Notice that ripples occur almost exclusively in the UP state. (Bottom) Rows in the pseudocolor image show the density of ripple occurrence for each dataset (n=87). Densities are replotted over two UDS cycles.

The distribution of DG CSD magnitude values is fit well by a binary Gaussian mixture (Figure 2C) consistent with synaptic activity switching between high and low level regimes. Since EC inputs are responsible for much of the dentate synaptic currents, we identify these regimes as corresponding to entorhinal UP and DOWN states (UDS). Since UDS in EC are coordinated with UDS in other cortical and thalamic areas, we reasoned that the DG CSD contains information about widespread brain state modulation and would therefore be correlated to other brain state signatures and behavioral metrics. Indeed we found that ripples tended to occur almost exclusively when the DG CSD was high (UP state), while eyeblinks had the opposite relationship and occurred when the DG CSD was low (DOWN state) (Figure 2C-E, Figure 2 - figure supplement 1). Furthermore, the DG CSD showed a strong coherence with the slow changes (below 1 Hz) in pupil diameter (Figure 2D, Figure 2 - figure supplement 1B).

In order to investigate these effects further we developed an unsupervised method for extracting UP and DOWN states from the DG CSD based on a hidden Markov model (HMM) (Figure 2C-D). In addition to identifying state transition points, the segmentation allows the translation of the time axis into a circular UDS phase, so that event distributions can be computed with respect to the UP-DOWN phase. This analysis demonstrates that essentially all ripples occur within the UP state and the rate of ripple occurrence ramps up to a steady state value in the course of the UP state itself and abruptly terminates upon transition to a DOWN state (Figure 2E). It also shows that upon transition to the DOWN state the pupil quickly dilates, signaling increased arousal, and then gradually constricts through the course of the UP state, indicating a progressive reduction of arousal and attention to external stimuli through the course of the UP state (Figure 2 - figure supplement 1D). Both UP and DOWN epoch durations were distributed approximately exponentially with means of 3.20 sec (UP) and 3.34 sec (DOWN) and these values were consistent across recording sessions (Figure 2 - figure supplement 2).

These observations show that quiet wakefulness can be reliably decomposed into UP and DOWN states that reflect increased and decreased EC inputs, respectively. This UDS classification reflects the dynamics of the cortical input to the hippocampus, in contrast to LIA/ SIA segmentation that is influenced by the states of the hippocampal subfields themselves, which as we describe below are not necessarily coherent.

### CA3 Membrane Potential Shifts are Negatively Correlated with Entorhinal Inputs to DG

What is the impact of entorhinal inputs on the activity of hippocampal neurons? The subthreshold membrane potential of the example CA3 neuron in Figure 2B is clearly related to the DG CSD and consequently to UP-DOWN states. Importantly, the relation is evident in the slow shifts in V_m_ and not in the superimposed faster fluctuations. This is confirmed by the fact that all significant coherence between V_m_ and DG CSD occurs below 1 Hz (Figure 2 - figure supplement 1B). We therefore separated the fast from the slow dynamics of the subthreshold membrane potential (V_m_) fluctuations with a cutoff frequency of approximately 1 Hz (Figure 2 - figure supplement 3).

The behavior of the example cell in Figure 2B is surprising because CA3 receives nearly identical excitatory ECII input as DG (Steward et al., 1976; Tamamaki and Nojyo, 1993), and yet the membrane potential of this CA3 pyramidal neuron is more hyperpolarized when DG CSD and hence the EC input rate is high and, conversely, more depolarized when EC input rate is low. Does EC input impact subthreshold activity in other hippocampal subfields in a similar way? To address this, we computed the cross-covariance between DG CSD magnitude and the slow V_m_ component of cells in DG, CA3, and CA1 (Figure 3). While the majority of DG and CA1 cells displayed a strong positive correlation to EC inputs, the majority of CA3 neurons exhibited a negative correlation. In other words, while the slow V_m_ fluctuations in DG and CA1 are nearly in sync with the DG CSD (Figure 3A, 3C), they are anticorrelated in CA3 (Figure 3B). Furthermore, a closer look at the lag associated with peak absolute correlation reveals that while DG neurons follow the DG CSD very closely (37 ms median lag), CA1 neurons actually lead the DG CSD (−227 ms median lag), while the trough of the CA3 negative correlation follows the DG CSD (296 ms median lag) (Figure 3 - figure supplement 1). This subfield ordering is inconsistent with a simple feedforward activation along the trisynaptic circuit in the awake state.

**Figure 3.**
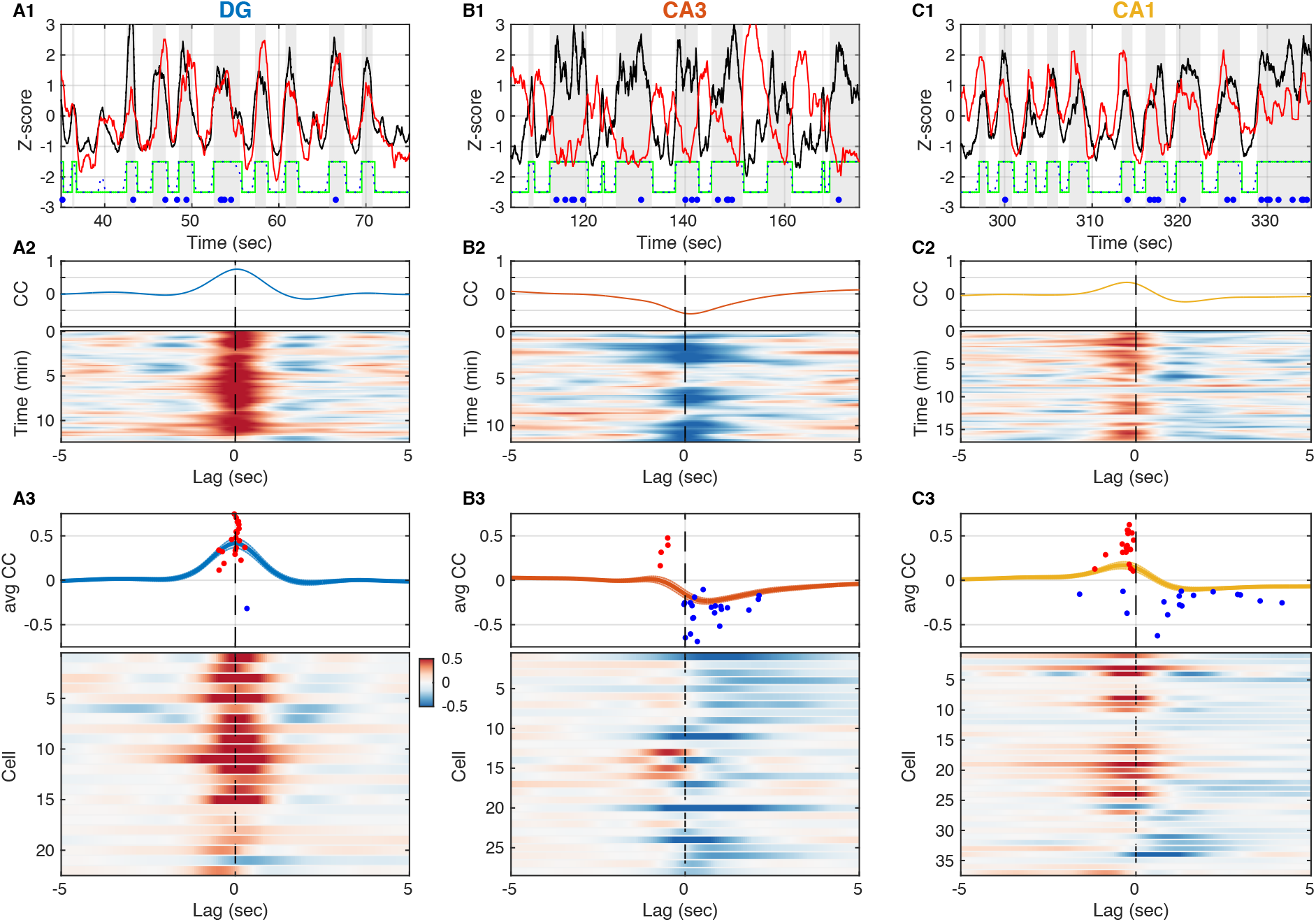
Entorhinal Input to DG is Negatively Correlated with Slow V_m_ Shifts in CA3 in Contrast to DG and CA1. **(A1)** DG CSD magnitude (black) and slow component (< 1 Hz) of subthreshold V_m_ (red) for an example DG granule cell. Gray vertical stripes and green stairstep trace mark periods classified as UP states. Ripples are marked by the blue dots. Notice that the V_m_ slow component is modulated in lockstep with the DG CSD. **(A2)** (Top) Cross-covariance between the slow V_m_ component of the example cell above and DG CSD magnitude. (Bottom) Cross-covariances are computed over 30 sec sliding windows and displayed as a pseudocolor image. **(A3)** (Top) Population average cross-covariance of all recorded DG granule cells. Dots mark the peak (red) or trough (blue) lag and amplitude of individual cells’ cross-covariance extrema. (Bottom) Cross-covariances for all DG granule cells stacked vertically and displayed as a pseudocolor image. Notice that most traces are peaked near zero lag. **(B)** Same as A, but for CA3 pyramidal neurons. Notice that the example cell in (B1, B2) and the population overall (B3) have membrane potentials that are anti-correlated with DG CSD magnitude. **(C)** Same as A and B, but for CA1 pyramidal neurons. The V_m_ of many CA1 neurons is positively correlated with DG CSD magnitude, but the V_m_ (red) leads the DG CSD (black) as in C1, so correlation peaks occur at negative lags (C3).

How well can we estimate the slow V_m_ component of hippocampal neurons from the DG CSD? All hippocampal subfields receive direct input from the entorhinal cortex (EC), and yet the pattern of modulation by the UP-DOWN state cycle varies across subfields. In order to understand how EC inputs influence hippocampal neurons we used the measured DG CSD as a proxy of EC input and estimated linear transfer models for each cell treating DG CSD as input and the cell’s slow V_m_ component as the model output (Figure 4). We considered a class of finite impulse response (FIR) models allowing for non-zero filter values at negative lags and hence for a non-causal influence of the input on the output. The estimated impulse responses (Figure 4A2-D2) revealed that while DG granule cells had causal responses, many CA3 and CA1 cells showed positive filter values at negative lags (up to -250 ms) signaling a non-causal relation between DG CSD and V_m_. This was consistent with the positive/negative peak correlation lags observed for DG/CA1 cells (Figure 3 - figure supplement 1A-B), but revealed a non-causal effect in CA3 which was not evident in the cross-covariance analysis. At positive lags the impulse responses for the majority of DG and CA1 cells were positive, while they were negative (inhibitory) for the majority of CA3 pyramidal neurons. The estimated models allowed us to simulate the step responses of hippocampal neurons (Figure 4A1-D1) and showed that as a population DG and CA3 exhibited persistent depolarization and hyperpolarization in response to sustained EC input, respectively, while CA1 showed a transient depolarizing response that decayed within 1 second. We also simulated the slow V_m_ components of hippocampal neurons from DG CSD and found good qualitative agreement with the experimental observations (Figure 4 - figure supplement 1), confirming that the transfer models provide a succinct description of the behavior of hippocampal neurons in response to changing EC input levels.

**Figure 4.**
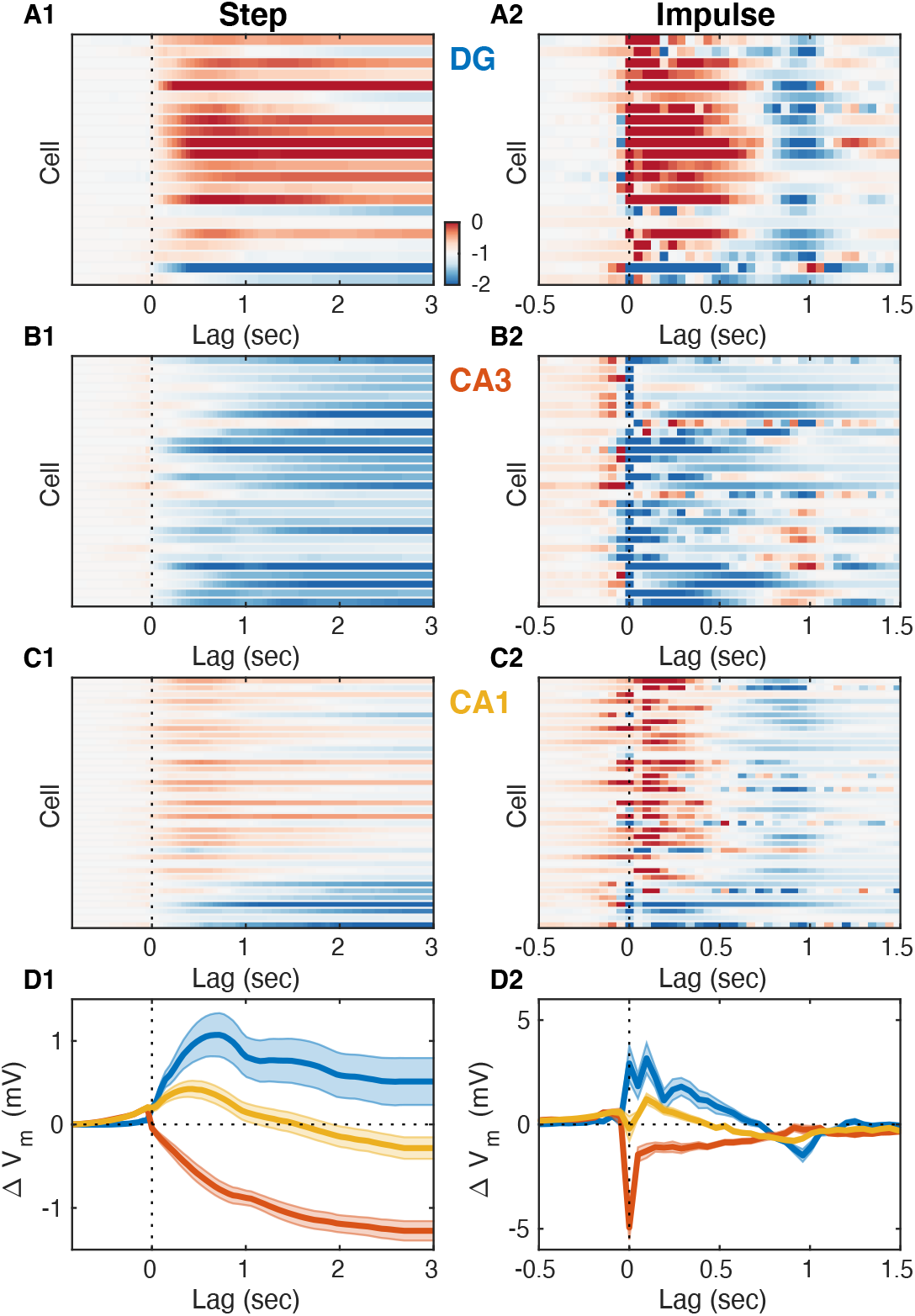
Entorhinal Input Differentially Modulates Slow V_m_ Shifts Across Hippocampal Subfields. **(A1)** Each row of the pseudocolor image shows the step response of a linear transfer model describing the effect of DG CSD magnitude on the slow V_m_ component for a given DG cell. The vertical interrupted line marks the onset of the input step. **(A2)** Each row shows the impulse responses of the corresponding models in A1. The vertical interrupted line marks the onset of the impulse. Notice that the majority of DG cells exhibit causal behavior, i.e. the impulse response is near zero for negative lags. **(B)** Same as A, but for CA3 pyramidal neurons. Notice that the majority of CA3 neurons hyperpolarize in response to EC input and some cells exhibit non-causal impulse responses, i.e. some impulse responses have non-zero (positive) values at negative lags. **(C)** Same as A and B, but for CA1 pyramidal neurons. Notice that some CA1 cells also exhibit non-causal impulse responses. **(D)** Area-specific population average step response (D1) and impulse response (D2) color-coded by brain area. Notice the distinct responses across hippocampal subfields.

### CA3 Neurons Hyperpolarize During UP States

The slow V_m_ component and firing rate of hippocampal neurons were modulated at UP and DOWN transitions (Figure 5 - figure supplement 1) in a way that was consistent with the transfer model predictions (Figure 5 - figure supplement 2). To understand their behavior through the course of the UDS cycle, we analyzed the subthreshold membrane potential and spiking of hippocampal neurons as a function of the UDS phase (Figure 5). In particular, we computed the distribution of V_m_ values conditioned on the phase of the UDS cycle. These distributions displayed phase-dependent V_m_ means for the majority of hippocampal neurons (Figure 5A1-D1) and many cells also exhibited phase-dependent fast V_m_ component variance (Figure 5A2-D2). Several important differences between the hippocampal subfields were evident. Dentate granule cells were quite homogeneous in their behavior through the course of the UP-DOWN state and all but one cell showed sustained V_m_ depolarization which was maintained through the course of the UP state and was mirrored by a persistent hyperpolarization throughout the DOWN phase (Figure 5A1). In contrast, about a third of CA3 neurons exhibited the exact opposite behavior: sustained V_m_ hyperpolarization through the UP state and depolarization through the DOWN state (Figure 5B1). The remaining population exhibited a more transient depolarization with a peak slightly preceding or coincident with the UP transition point (Figure 5B1). The CA1 pyramidal cell population displayed a level of diversity that was intermediate to that of DG and CA3. About two thirds of CA1 neurons were depolarized in the UP state and hyperpolarized in the DOWN state, but the responses were more transient than in DG, with depolarization/hyperpolarization decaying through the course of the UP/DOWN state, respectively (Figure 5C1). The behavior of the remaining CA1 neurons appeared similar to that of the CA3 population. With respect to V_m_ fluctuations, DG granule cells exhibited the highest variability, exceeding that in CA3 and CA1 (Figure 5 -figure supplement 3A1-D1). With respect to overall V_m_ fluctuations only CA1 neurons displayed state-dependent modulation of V_m_ variability (Figure 5 - figure supplement 3D1), with the UP state being associated with more variable V_m_. Focusing on the fast V_m_ component alone, both CA1 neurons and DG granule cells showed pronounced jump in variability during the UP state (Figure 5A2-D2). In contrast, CA3 pyramidal neurons exhibited a nearly constant level of V_m_ variability throughout the UP-DOWN state cycle, which equaled that of CA1 in the UP state, but exceeded it in the DOWN phase (Figure 5B2, 5D2, Figure 5 - figure supplement 3D1). Interestingly, with respect to spiking the population of DG and CA1 neurons exhibited similar behavior, a sustained increase in firing rate that persisted through the course of the UP phase (Figure 5A3-D3). In contrast, the firing behavior of CA3 neurons largely mirrored the behavior of the V_m_ mean, with about a third of CA3 neurons exhibiting a sustained increase in firing rate throughout the course of the DOWN state and the remaining population showing a transient increase peaking before or at the UP transition point (Figure 5B3, 5D3). Thus CA3 neurons were maximally hyperpolarized and had lowest firing probability at the end of the UP phase when the rate of ripple occurrence was highest (Figure 2E).

**Figure 5.**
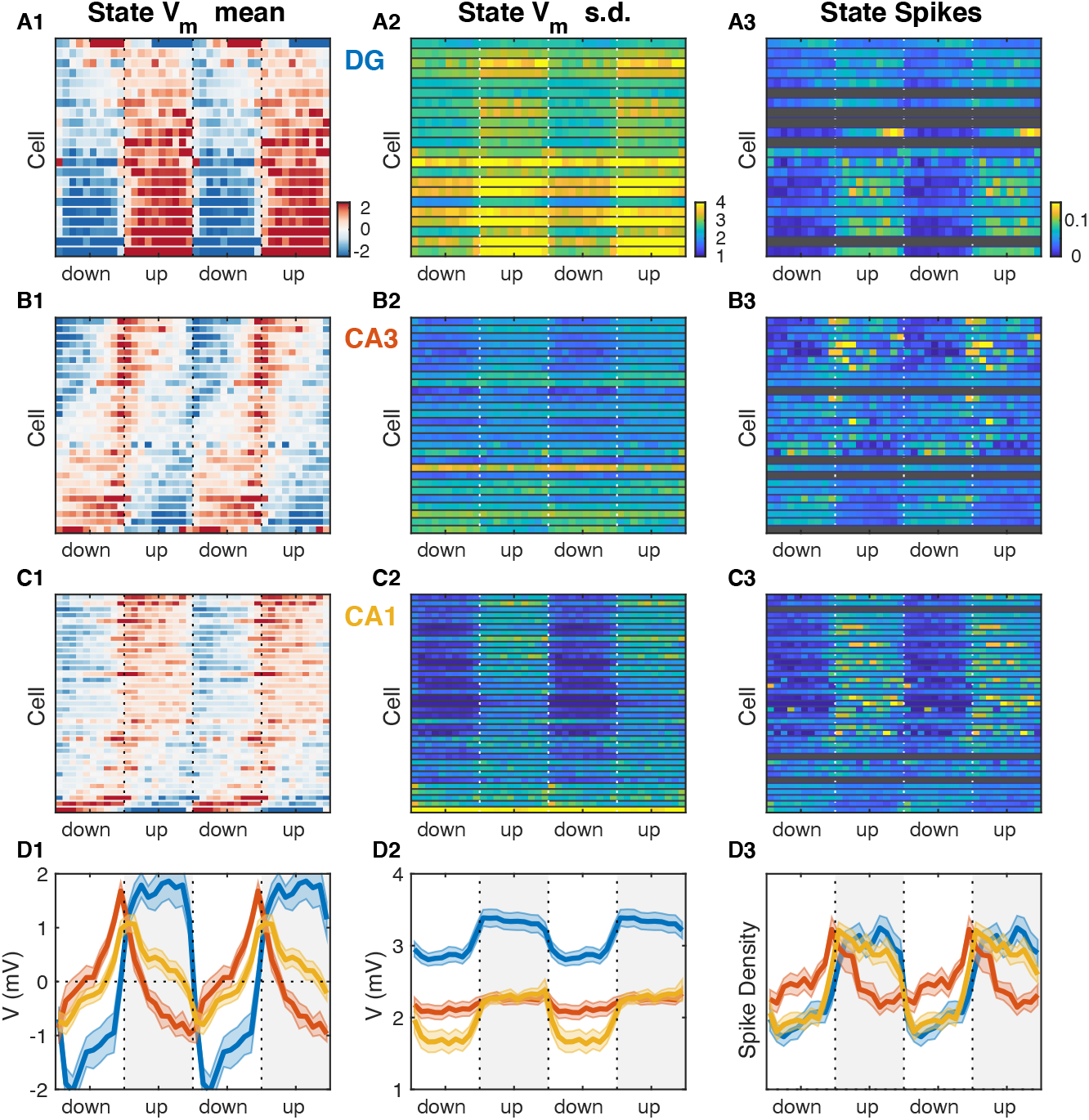
Hippocampal Subfields Exhibit Distinct Activity Profiles over the UDS Cycle. **(A1)** V_m_ mean as a function of UDS phase for each DG granule cell displayed as a row in the pseudocolor image. **(A2)** Fast V_m_ component standard deviation as a function of UDS phase for DG granule cells. **(A3)** Observed spiking probability for all DG granule cells displayed as a pseudocolor image. Grayed out rows correspond to cells that fired fewer than 100 spikes. **(B)** Same as A, but for CA3 pyramidal neurons. Notice that CA3 neurons are maximally hyperpolarized and have lowest firing probability at the end of the UP phase when the rate of ripple occurrence is highest (Figure 2E). **(C)** Same as A and B, but for CA1 pyramidal neurons. **(D)** Area-specific population averages and examples color-coded by brain area. (D1) Population average V_m_ means. (D2) Population average fast V_m_ component standard deviations. (D3) Population average spiking probability.

### Inhibition Dominates CA3 Subthreshold Behavior near Awake Ripples

It has long been appreciated that awake ripples preferentially occur during periods of quiet wakefulness and LIA. Furthermore, here we showed that when quiet wakefulness is segmented into periods of UP and DOWN states, essentially all ripples occur in the EC UP state (Figure 2E). The prevailing view is that ripples are the result of self-organized population bursts of activity that build within the recurrent CA3 network and then are transmitted to and further patterned by the CA1 network, where the ripple oscillation itself is readily observable in the local field near the pyramidal cell layer (Figure 2A) (Buzsáki, 1986; Miles and Wong, 1983; Stark et al., 2014; Traub and Miles, 1991). How do neurons across the different hippocampal subfields participate in this process? To address this question we detected ripple onset times from the CA1 local field near the pyramidal cell layer and determined the ripple-triggered average (RTA) V_m_ and firing rate modulation for each recorded cell (Figure 6). Since the RTA of the V_m_ is the sum of the RTAs of the fast and slow V_m_ components, we determined these average responses separately for each cell and component (Figure 6A1-D1, 6A3-D3). The RTAs of the slow V_m_ component showed pronounced depolarization in DG with a peak before the ripple onset, a hyperpolarization in CA3 with a trough following the ripple onset, and mixed responses in CA1 (Figure 6A1-D1). Since ripples occurred exclusively during the UP state and the slow V_m_ component of hippocampal neurons was strongly influenced by state, we hypothesized that the RTA slow V_m_ response depended strongly on the pattern of EC input and how each cell was impacted by it. We therefore simulated the slow V_m_ component for each cell using the linear transfer models we had estimated (Figure 4) and computed the ripple-triggered average of the simulated slow V_m_ response (Figure 6A2-D2). The RTAs of the observed and simulated slow V_m_ components showed good qualitative agreement indicating that on the timescale of seconds V_m_ trends near ripples are strongly influenced by the pattern of EC UDS transitions.

**Figure 6.**
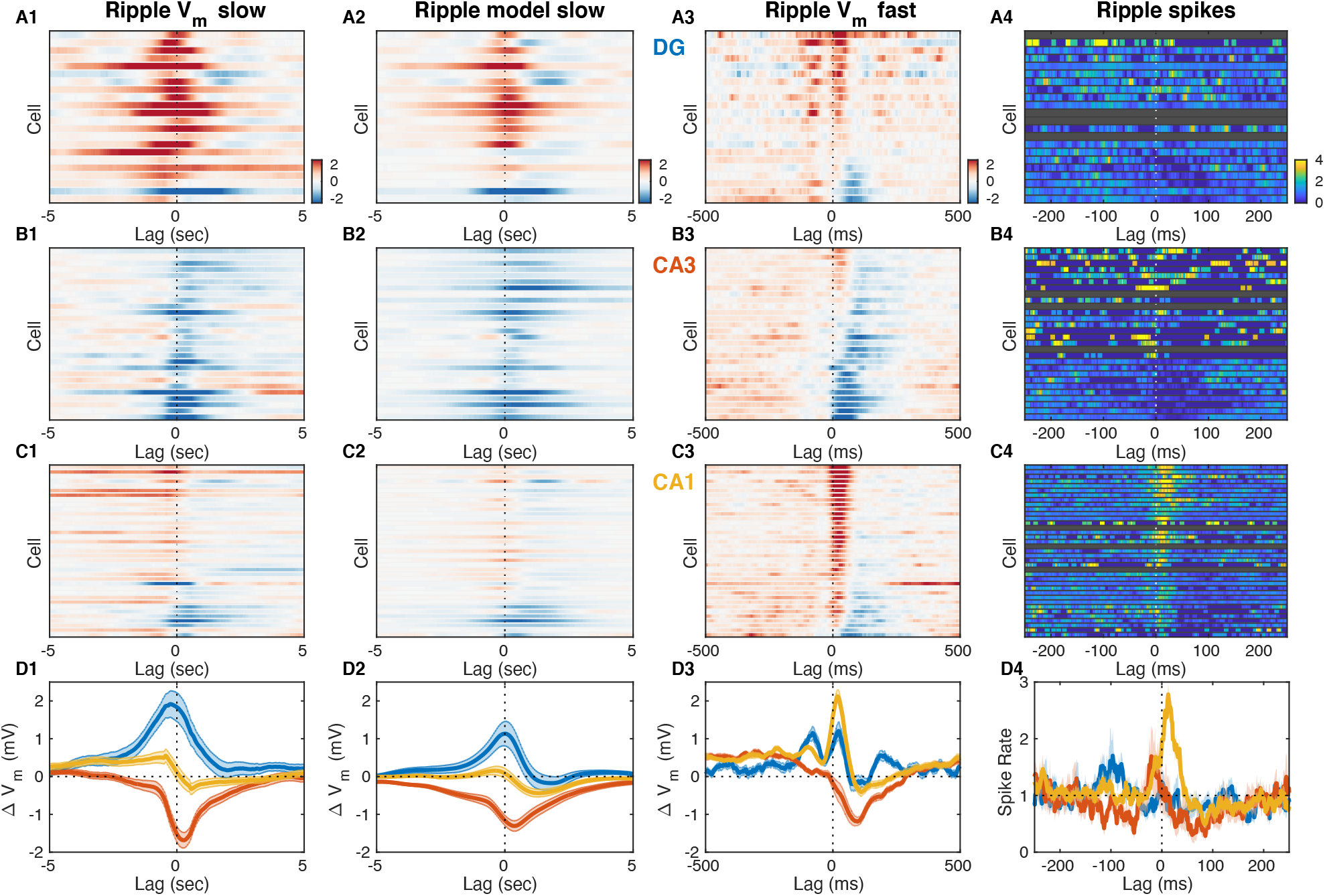
Inhibition Marks Slow and Fast V_m_ Responses near Ripples in CA3, Unlike DG or CA1. **(A1)** Mean slow V_m_ component triggered on ripple onset for each DG granule cell displayed as a row in the pseudocolor image. Interrupted vertical line marks ripple onset. **(A2)** Transfer model predicted ripple-triggered slow V_m_ component as in A1. **(A3)** Mean fast V_m_ components triggered on ripple onset and displayed as in A1. Notice the two depolarizing peaks at -100 ms and right after ripple onset. **(A4)** Spiking response to ripple onset for DG cells normalized by baseline firing rate. **(B)** Same as A, but for CA3 pyramidal neurons. Notice the prominent hyperpolarization for most CA3 neurons present in both the slow and fast V_m_ component responses. **(C)** Same as A and B, but for CA1 pyramidal neurons. Notice the prominent depolarization for most CA1 neurons. **(D)** Area-specific population average slow V_m_ response (D1), transfer model predicted slow V_m_ responses (D2), fast V_m_ response (D3), and spiking response (D4) color-coded by brain area. Notice that modulation by UDS accounts for the shape of the ripple-triggered V_m_ response on the timescale of seconds (compare D1 and D2).

In contrast, the mean of the fast V_m_ component is very weakly influenced by state and therefore its RTA should reflect synaptic activity consistently timed with respect to the ripple onset. Consistent with previous studies, we found that the fast RTA V_m_ waveform for most CA1 pyramidal neurons had a sharp prominent peak following the ripple onset, followed by a steep return to baseline or hyperpolarization within a 100 ms (Figure 6C3) (Hulse et al., 2016). This behavior was mirrored in the firing rate modulation of CA1 cells (Figure 6D3) and is consistent with the notion that CA1 exhibits a highly synchronous population burst associated with ripple oscillations.

Since the population event in CA1 is presumably due to a self-organized population burst in CA3, one might expect that the fast RTA waveforms in CA3 should resemble those in CA1, but this was not the case (Figure 6B3). While a minority CA3 neurons exhibited a small depolarizing peak near the ripple-onset, the most consistent feature of the fast RTA V_m_ waveform in CA3 was the prominent hyperpolarization that reached a trough roughly 100 ms following ripple onset and recovered within 300 ms. The population average response shows that the decrease in V_m_ had begun as early as 150 ms prior to the ripple onset (Figure 6D3). Similarly, while several CA3 neurons showed an increase in firing rate just prior to the ripple onset, about a third of the CA3 population exhibited a reduction in firing rate before and especially following the ripple onset (Figure 6B4, 6D4). These data demonstrate that extensive inhibition, rather than excitation, is the hallmark of CA3 subthreshold activity before and after ripple bursts. It is worth noting that most neurons, even with hyperpolarizing average response, experienced depolarization and firing during a subset of ripples, consistent with a sparse activation of CA3 during ripples.

Finally, while the dentate gyrus has not been traditionally considered to be instrumental to the process of ripple generation, the fast RTA V_m_ waveforms of granule cells showed clear modulation with about half of the population exhibiting a depolarization peak following ripple onset with a similar time course to that seen in CA1 (Figure 6A3). Interestingly, about half of DG granule cells also showed another depolarization peak occurring 100 ms prior to ripple onset (Figure 6A3, 6D3). Upon closer examination a similarly timed depolarizing peak could be seen in the CA1 response prior to ripple onset, although of much smaller magnitude in comparison to the post-ripple depolarization. Similarly, a sharp wave of smaller amplitude is also consistently observed 100 ms before ripple onset (Hulse et al., 2016). These data indicate that DG should not be viewed as a passive bystander in the awake ripple generation process and that the ripple proper is preceded by coordinated activity not only in CA3, but also in CA1 and DG.

What circuit mechanisms may be responsible for the observed subthreshold activity around ripples? Since CA3 is considered to play a key role in the process of ripple generation, we reasoned that influences on subthreshold activity originating in CA3 should scale in proportion to the CA3 population burst size. We inferred this size indirectly from the amplitude of the ripple-associated sharp wave in stratum radiatum of CA1 or equivalently by the magnitude of the underlying synaptic current due to Schaffer collateral activation (Mizunuma et al., 2014). We then divided the ripples from each recording session in two halves, big and small, depending on whether they were associated with above-median or below-median CA3 burst size. Finally, we computed the fast RTA V_m_ waveform separately for big and small CA3 events and compared the results (Figure 7). The size of the CA3 burst had little effect on the DG RTA waveform, apart from a slight increase in hyperpolarization about 100 ms following ripple onset that was associated with the bigger events (Figure 7A1-A3). In contrast, subthreshold activity in CA3 clearly reflected the size of the CA3 population event (Figure 7B1-B3). Larger CA3 events were associated with more sustained depolarization up to 300 ms prior to the ripple onset and deeper hyperpolarization following (Figure 7B3). In CA1, bigger CA3 events were associated with slightly more depolarization up to 250 ms prior to the ripple onset and during the post-ripple peak, but the most notable difference was the increased hyperpolarization about 100 ms following the ripple onset (Figure 7C1-C3). These data indicate that feedback inhibition within CA3 is most likely responsible for the subthreshold hyperpolarization seen in the fast V_m_ component around ripples in CA3. Consistent with our previous study, it also suggests that feedforward inhibition from CA3 to CA1 may play an important role in controlling the population event size in CA1 and may contribute to the post-ripple hyperpolarization (Hulse et al., 2016).

**Figure 7.**
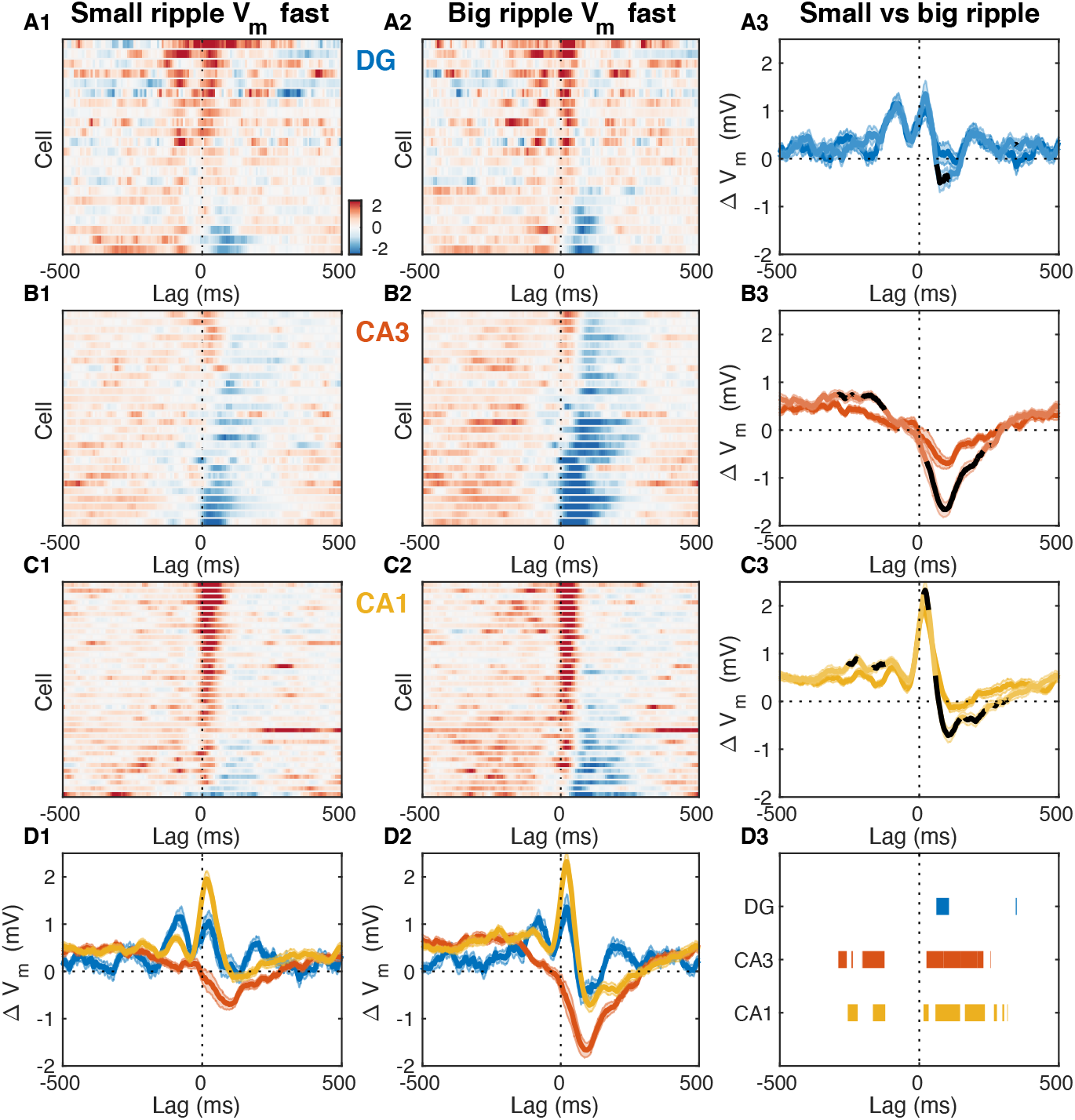
Fast V_m_ Inhibitory Responses to Ripples Scale with CA3 Population Burst Size. **(A1)** Mean fast V_m_ components of DG granule cells triggered on ripples with small (below average) sharp-wave (SPW) amplitudes. **(A2)** Same as A1, but for ripples with big (above average) SPW amplitudes. **(A3)** Comparison of the DG population average response to small and big SPW ripples. **(A4)** Significant differences are highlighted in black. **(B)** Same as A, but for CA3 pyramidal neurons. **(C)** Same as A and B, but for CA1 pyramidal neurons. **(D)** Area-specific population average responses to small (D1) and big (D2) SPW ripples. (D3) Horizontal bars mark the onset and duration of significant differences between responses to small and big SPW ripples within each area.

## DISCUSSION

The results above show that the V_m_ modulation of CA3 pyramidal neurons by entorhinal input and ripples is largely opposite to that of DG granule cells and CA1 pyramidal neurons. On the timescale of seconds, slow shifts in the V_m_ of CA3 neurons are negatively correlated to the UP-DOWN transitions in the level of entorhinal input to DG. Consequently many CA3 neurons exhibit hyperpolarization in the UP state in contrast to DG and CA1 neurons, which are in sync with EC inputs. This subfield-specific modulation by UDS explains the slow trends in the membrane potential leading to and following awake ripples. On a sub-second timescale, both DG and CA1 neurons exhibit depolarization transients both ∼100 ms before and immediately after the ripple onset, while CA3 neurons show a prominent hyperpolarization that starts to build before and reaches maximum after the ripple onset. The magnitude of the CA3 hyperpolarization scales with the size of the CA3 population burst pointing to feedback inhibition as its likely source.

### UDS Differentially Modulate Activity Across DG, CA3, and CA1

Neocortical dynamics during NREM sleep and anesthesia show intrinsic alternation between periods of elevated activity (UP states) and relative quiescence (DOWN states) (Cowan and Wilson, 1994; Steriade et al., 1993a). The entorhinal cortex is the major gateway linking the neocortex with the hippocampal formation and has been shown to exhibit UP and DOWN states (UDS) associated with bimodal membrane potential distributions of EC neurons (Isomura et al., 2006). Despite an absence of bimodality, the subthreshold activity and the spiking of hippocampal neurons is modulated by cortical UDS in a subfield-specific manner both in sleep and under anesthesia (Hahn et al., 2007; Isomura et al., 2006; Sullivan et al., 2011). We found that DG molecular layer currents exhibit UP-DOWN dynamics indicating that the EC undergoes UDS transitions in quiet wakefulness as well that modulate activity across hippocampal subfields.

Our results regarding UDS modulation in quiet wakefulness are largely consistent with corresponding observations in NREM sleep and under anesthesia, but there were also some notable differences that we highlight below. We observed pronounced modulation of both subthreshold activity and spiking at the DOWN→UP transition in all hippocampal subfields including a 50% increase in the baseline firing rate of CA3 neurons preceding the DOWN→UP transition in contrast to previous reports which found no modulation of CA3 unit activity in sleep/waking immobility (Sullivan et al., 2011) or weak and mixed modulation in anesthetized animals (Hahn et al., 2007). We also find that awake ripples occur essentially exclusively during the UP state, and not merely with an increased probability relative to the DOWN state (Sullivan et al., 2011). Furthermore, awake ripples occurred throughout the UP state and were not concentrated near the DOWN→UP transitions (Battaglia et al., 2004). Finally, in our data the vast majority of CA1 neurons were depolarized on the DOWN→UP transition and had elevated firing rates in the UP state in contrast to previous observations under anesthesia (Hahn et al., 2007; Isomura et al., 2006).

While the majority of recorded neurons were influenced by UDS, the nature of the modulation is subfield-specific. Granule cells in DG by and large follow the EC inputs and show sustained depolarization and firing rate increase throughout the UP state mirrored by relative hyperpolarization and reduced firing in the DOWN state. Although CA3 follows DG in the trisynaptic loop, DG activity is more similar to that in CA1 than in CA3. In particular, CA1 pyramidal neurons also depolarize and fire more on the DOWN→UP transition, but these responses are more transient than in DG. Consequently, the expected membrane potential across the CA1 population has a triangular wave shape as a function of UDS phase, unlike the square wave shape characteristic of the DG population. This triangular wave shape is almost symmetric with respect to DOWN→UP transition and as a result the V_m_ conditional means in the UP and DOWN states are very similar, despite the clear V_m_ modulation by UDS phase. The CA1 population starts depolarizing before the DOWN→UP transition and the DG granule cells. This is reflected in the non-causal transfer model impulse response of CA1 pyramidal cells, which is consistent with the presence of a feedback loop via the CA1→EC connection, but also suggests the presence of another excitatory source, such as CA3, that leads EC activity.

How can activity in CA3 lead given that CA3 is downstream of both the EC and the DG? The majority of CA3 pyramidal neurons are negatively correlated to DG molecular layer currents, which is surprising since they receive essentially the same excitatory input from layer 2 of the EC as DG granule cells, while the mossy fibers (DG→CA3) form powerful excitatory “detonator” synapses on the proximal dendrites of CA3 pyramidal neurons. Despite this anatomical organization suggesting that CA3 activity should follow that in EC and DG, CA3 pyramidal cells in fact show peak depolarization and elevated firing before the DOWN→UP transition, thus leading both CA1 and DG. Furthermore, a third of the CA3 population is more depolarized in the DOWN state, while the rest exhibit transient depolarization right before the DOWN→UP transition. This is reflected in the negative impulse response of the CA3 transfer model with sustained negative step response. The modulation of the population firing rate in CA3 by UDS is consistent with CA3 being responsible for the CA1 lead over DG activity.

What mechanisms may account for the CA3 behavior? One possibility is that DG and/or EC inputs provide powerful feedforward inhibition to CA3 pyramidal neurons. Indeed mossy fibers (DG→CA3) not only form large mossy terminals on CA3 pyramidal cells but also contact interneurons via filopodial extensions, providing an anatomical substrate for feedforward inhibition (Acsády et al., 1998). The balance between feedforward excitation and inhibition depends on the pattern of granule cell activity: low frequency activation of the the mossy fibers results in powerful slow inhibition of CA3 pyramidal neurons while at higher frequencies an initial depolarization precedes the inhibition (Zucca et al., 2017). Through this mechanism the elevated DG activity in the UP state may induce a concomitant suppression in the majority of CA3 pyramidal neurons during the UP state. As granule cells reduce their firing in the DOWN state, the CA3 circuit is released from the DG-mediated feedforward inhibition and the recurrent CA3 connections may support a sustained increase in population activity. This recurrent excitation is controlled by the strong feedback inhibition present in CA3. The non-causal positive component of the CA3 transfer model impulse response and the timing of CA3 firing relative to the DOWN→UP transition indicate that CA3 may play an important role in ushering the subsequent UP state by providing excitation to EC via CA1. These observations are inconsistent with a feedforward activation of the trisynaptic pathway, suggesting a more complex interplay of intrahippocampal and perforant pathways.

### Membrane Potential Dynamics Around Ripples in Quiet Wakefulness

The recurrent circuit of CA3 has long been hypothesized to function as an autoassociative memory network (Marr, 1971) and to support the buildup of population activity underlying the ripple generation process (Buzsáki, 2015). Indeed, acute silencing of Schaffer collaterals during wakefulness abolishes ripples (Davoudi and Foster, 2019; Yamamoto and Tonegawa, 2017). While the membrane potential of CA1 pyramidal neurons near ripples has been shown to exhibit a gradual ramping and a transient depolarization followed by a prolonged inhibition (Hulse et al., 2016), the subthreshold dynamics of DG granule cells and CA3 pyramidal neurons near ripples in awake animals had not been fully characterized.

We observed that the ripple-triggered membrane potential of hippocampal neurons is modulated on a timescale of seconds, with DG granule cells showing depolarization, CA3 pyramidal neurons hyperpolarization, and CA1 cells exhibiting weaker modulation. Since ripples occur almost exclusively in the UP state and the slow V_m_ dynamics of hippocampal cells are modulated by UDS in a subfield-specific fashion, we hypothesized and confirmed that the slow V_m_ responses near ripples can be qualitatively accounted for by the UDS influence on hippocampal cells.

How does UDS influence hippocampal network excitability? It has been proposed that during NREM sleep hippocampal dynamics exhibit a stable, but excitable quiescent state, such that activity fluctuations can produce a transient population excitation representing a ripple (Levenstein et al., 2019). Our data indicate that in quiet wakefulness UDS modify hippocampal network excitability because no ripples are produced in the DOWN state. We illustrate this behavior in a mean firing rate model in the framework described in (Levenstein et al., 2019), featuring two (EC and CA3) adapting recurrent neural populations (Figure 8). In the model the EC population influences CA3 activity by providing a net inhibitory input as well as by modulating the strength of the CA3 recurrent excitation (Figure 8A). In the real circuit the latter influence may be due to UDS-dependent changes in neuromodulatory inputs, such as cholinergic tone, as exhibited, for example, by pedunculopontine cholinergic neurons (Mena-Segovia et al., 2008). The majority of the cholinergic input to the hippocampus originates in the medial septum and optogenetic stimulation of septal ChAT-positive neurons suppresses ripple generation (Hunt et al., 2018; Vandecasteele et al., 2014). Acetylcholine is known to inhibit the efficacy of recurrent synaptic transmission in CA3 by acting on presynaptic muscarinic receptors in the associational–commissural fiber system (Hasselmo et al., 1995; Hasselmo and Schnell, 1994; Vogt and Regehr, 2001) and cholinergic tone is presumably at its lowest during ripple generation in the EC UP state. Thus the efficacy of CA3 recurrent connections together with dentate and entorhinal input to CA3 and the associated feedforward inhibition can act as bifurcation parameters for the CA3 network dynamics that change network excitability thereby preventing ripple occurrence during the cortical DOWN state (Figure 8E-F). This is counterintuitive because many CA3 neurons are more active and depolarized in the DOWN state or near the DOWN→UP transition and so, according to the stochastic-refractory model of ripple initiation (Schlingloff et al., 2014) the rate of ripple occurrence should coincide with the modulation of CA3 activity by UDS, which is contradicted by the fact that ripples occur in the UP state. In our model, CA3 network excitability is shown to be restricted to the EC UP state when the mean CA3 rate is lower compared to the DOWN state (Figure 8B-C). This is possible because of the push-pull influences of the net inhibitory input which lowers CA3 network excitability, and the CA3 recurrent strength potentiation which increases it. Hence, in the UP state the CA3 network is inhibited but excitable, while in the DOWN state it is disinhibited but not excitable (Figure 8).

**Figure 8.**
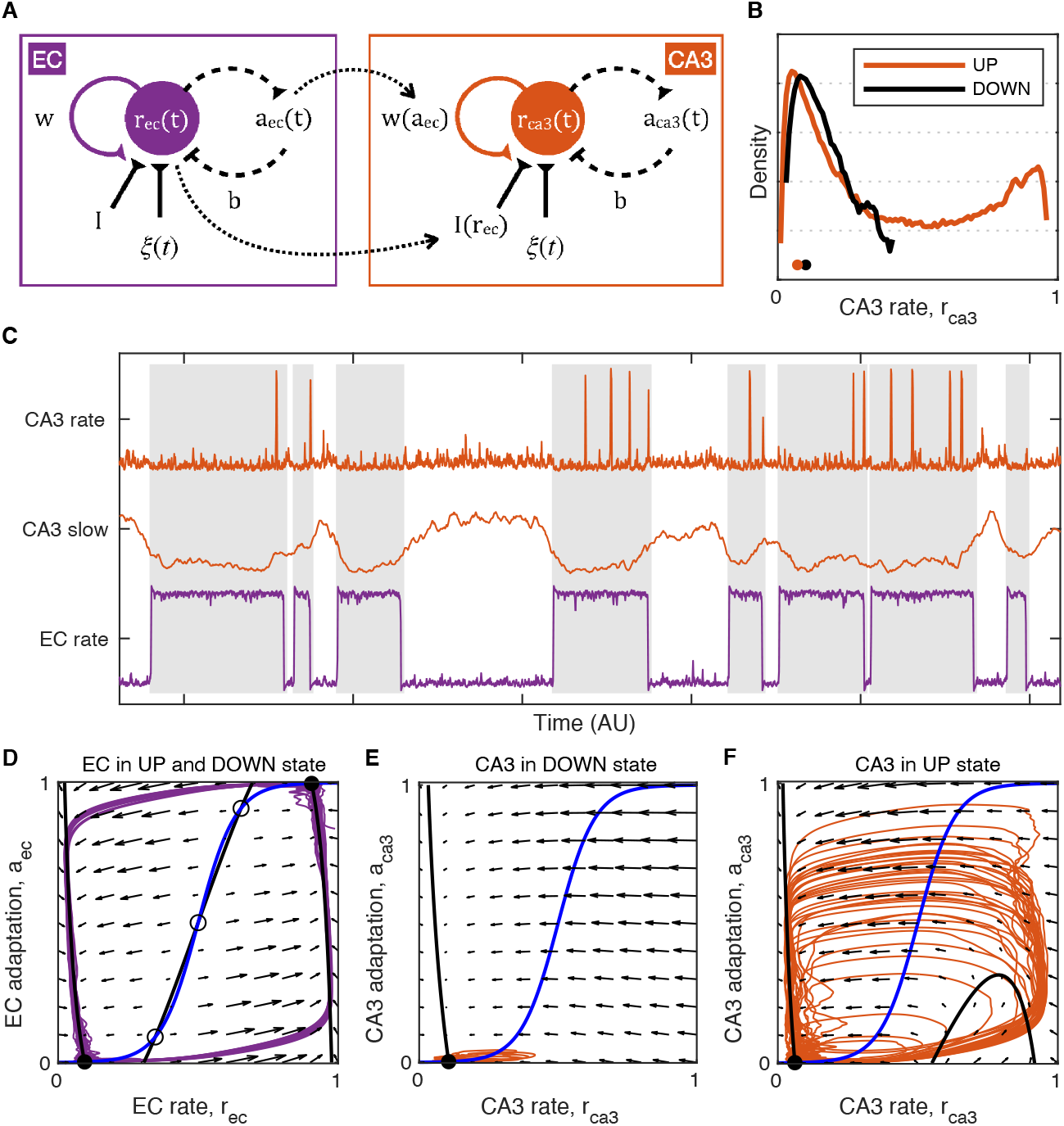
Model of UDS Control of CA3 Network Excitability. **(A)** Idealized model of EC and CA3 adapting recurrent neural populations. EC activity provides net inhibition to the CA3 population via *I*(*r*_*ec*_), possibly due to feedforward inhibition or indirect influence via DG, and modulates the strength of CA3 recurrent excitation via *w*(*a*_*ac*_), modeling the effects of UDS-dependent shifts in cholinergic tone on CA3 synaptic transmission. **(B)** Probability density of the simulated CA3 population rate r *ca*3 during EC UP and DOWN states. Notice that the median CA3 population rate is higher in the DOWN than the UP state (black and orange dots) despite the presence of population bursts (rate values near 1) restricted only to the UP state. **(C)** Example model simulation demonstrating that the EC population exhibits UP and DOWN dynamics while the CA3 population produces transient population bursts (“ripples”) restricted to the UP state (gray segments). The slow component of the CA3 rate is magnified in the middle to show that mean CA3 activity is higher during the DOWN state and lower during the UP state when population bursts occur. **(D-F)**Phase plane plots of the model dynamics. The model evolution is governed by the velocity fields displayed as arrows. The model stable (filled circles) and unstable (open circles) fixed points occur at the intersections of the population rate (black) and adaptation (blue) nullclines. Model trajectories are plotted in purple and orange. **(D)** EC dynamics exhibit two stable fixed points (black circles) corresponding to the UP and DOWN states with noise fluctuations driving transitions between them. **(E)** CA3 dynamics in the EC DOWN state exhibit a single stable fixed point (black circle) and are not excitable, i.e. noise fluctuations cannot trigger a spike in the population rate. **(F)** CA3 dynamics in the EC UP state exhibit a single stable fixed point (black circle) at a lower population rate level than in the DOWN state (E), but are excitable, i.e. noise fluctuations can trigger population spikes (“ripples”).

These data suggest that increased inhibition in CA3 may be a prerequisite for ripple initiation. This is consistent with *in vitro* work showing that reducing GABA_A_-mediated inhibition in hippocampal slices abolishes spontaneously occurring sharp wave-ripple events in CA3 (Bazelot et al., 2016; Ellender et al., 2010; Schlingloff et al., 2014). The relative hyperpolarization in CA3 during the UP state may reflect the role of certain local (Katona et al., 2014; Viney et al., 2013) or long-range projecting interneurons (Basu et al., 2016; Unal et al., 2018) in suppressing the initiation of ripples. According to one view, ripple generation may be the result of disinhibition, however our data does not offer clear evidence for widespread disinhibition in CA3 preceding the population burst (Evangelista et al., 2020). Importantly, ripple generation by disinhibition does not account for the lack of ripples in the DOWN state, despite the elevated neuronal activity in CA3. Its origin notwithstanding, membrane hyperpolarization in CA3 may influence ripple initiation by affecting voltage-gated conductances and thereby changing the excitability of CA3 neurons to make them more likely to fire or burst in response to transient depolarizing input.

On a faster timescale, the average V_m_ responses near ripples reveal two consistent features. First, the majority of CA3 neurons exhibit a brief (∼300 ms) hyperpolarization locked to the ripple onset in addition to the broad (∼3 sec) UDS-mediated hyperpolarization. This brief hyperpolarization grows with the size of the CA3 population burst, quantified by the associated sharp wave amplitude, therefore pointing to feedback inhibition as its source. This suggests that feedback inhibition is a consistent feature of the buildup process in CA3. This inhibition likely arises from several classes of interneurons that have been shown to exhibit elevated firing around ripples in both CA3 and CA1 (Klausberger et al., 2003; Somogyi et al., 2014; Tukker et al., 2013). However, the distinct V_m_ responses we observed around ripples indicate that inhibition is tuned differently in CA3 and CA1. In particular, in CA1 inhibition imposes oscillatory patterning on the V_m_ that rides on a wave of depolarization, while in CA3 inhibition summates to produce a net hyperpolarization. In both areas however fast fluctuations in the membrane potential persist throughout the ripple period allowing CA3 neurons to fire despite the net hyperpolarization. These results suggest that the relative gain of feedback inhibition is greater than that of recurrent excitation for the majority of the CA3 neurons and hence the population burst can only build up by recruiting the CA3 neurons most strongly connected to a sparse active subset. This may reflect a winner-take-all mechanism for controlling both the sparsity and the specificity of the neuronal sequences activated in CA3 during a ripple. The growing inhibition during the course of a ripple also provides a mechanism for ripple termination.

Second, DG granule cells exhibit two transient depolarizations of comparable amplitude ∼100 ms before and immediately after the ripple onset. Similarly timed features are present in the V_m_ of CA1 pyramidal neurons, albeit the pre-ripple depolarization is significantly smaller than the post-ripple one. In CA1, both depolarizations are associated with sharp waves of proportional magnitudes in stratum radiatum (Figure 4A in (Hulse et al., 2016)), implicating CA3 as the source for both. The DG activation is consistent with the presence of a backprojection from CA3 to DG (Scharfman, 2007; Szabo et al., 2017) and can influence ripple-associated CA3 activity, consistent with previous lesion studies (Sasaki et al., 2018). These results indicate a long and orchestrated ripple initiation process in the awake state, extending beyond the roughly 50 ms period of excitatory activity buildup that proceeds ripples *in vitro* (Schlingloff et al., 2014).

These results provide novel insights into the processes of ripple initiation, build up, and termination in awake animals. Ripples occur exclusively in UP states characterized by increased entorhinal inputs to DG and associated with pronounced hyperpolarization of CA3 pyramidal cells. This suggests that broad inhibition in CA3 may be a prerequisite for ripple initiation. Furthermore, DG and CA1 pre-ripple responses suggest that ripples are not initiated as isolated events within CA3, but are the culmination of extended interplay across multiple areas. This may reflect the role of cortical inputs in influencing the neuronal patterns replayed by the hippocampus during awake ripples, consistent with their role in spatial decision making. Finally, growing hyperpolarization in CA3 throughout the course of a ripple suggests that feedback inhibition is a key feature of ripple buildup as well as termination. This may reflect a winner-take-all mechanism, by which a few neurons that fire overcoming a background of inhibition in UP states, further suppress all other neurons via feedback inhibition, ensuring sparseness and selectivity of transient network pattern activation.

## MATERIALS AND METHODS

### Head fixation surgery

The methods used were the same as those described in our previous publications (Hulse et al., 2017, 2016). Briefly, male C57Bl/6 mice (Charles River Laboratories) were surgically implanted with a light-weight, stainless steel ring using dental cement. A stainless steel reference wire was implanted over the cerebellum for LFP silicon probe recordings. The locations of future craniotomies for probe and whole-cell recordings were marked. Probe recording coordinates were anterioposterior (AP)/mediolateral (ML): -1.7/1.75 in the left hemisphere for DG; AP/ML: -1.7/2.0 in the left hemisphere for CA1; and AP/ML: -2.15/0.84 in the right hemisphere for CA3. Whole-cell recording coordinates were: AP/ML: -1.7/0.65 in the left hemisphere for DG; AP/ML: -1.9/1.5 in the left hemisphere for CA1; and AP/ML: -2.15/3.1 in the right hemisphere for CA3. All coordinates are reported in mm, all AP and ML coordinates are with respect to bregma. Following surgery, mice were returned to their home cage, maintained on a 12 hour light/dark cycle, and given access to food and water ad libitum. Ibuprofen (0.2 mg/mL) was added to the water as a long-term analgesic. Mice were given at least 48 hours to recover before the day of the experiment.

### Exposure surgery

On the day of the recording, while mice (4-8 weeks old) were anesthetized with 1% isoflurane and head-fixed in the stereotaxic apparatus, two small craniotomies (∼200-500 µm) were made at the previously marked locations. A recording chamber was secured on top of the head-fixation device and filled with pre-oxygenated (95% O2, 5% CO2), filtered (0.22 μm) artificial cerebrospinal fluid (aCSF) containing (in mM): 125 NaCl, 26.2 NaHCO3, 10 Dextrose, 2.5 KCl, 2.5 CaCl2, 1.3 MgSO4, 1.0 NaH2PO4.

### Awake, *in vivo* recordings

Awake, *in vivo* electrophysiological recordings were carried out following previously described methods (Hulse et al., 2017, 2016). Mice were head-fixed on a spherical treadmill secured on an air table (TMC). A single-shank, 32-site silicon probe (NeuroNexus) with 100 μm site spacing was inserted in the coronal plane (∼15 degree angle pointing towards the midline) to a depth of 2600-3400 μm and was adjusted for reliably recording LFP ripple oscillations in CA1. To find the rough target depth of whole-cell recording, at first juxtacellular recordings (Pinault, 1996) were performed with pipettes filled with artificial cerebrospinal spinal fluid (aCSF). Whole-cell patch-clamp recordings were performed with a blind-patch approach (Margrie et al., 2002; Pinault, 1996) in current clamp mode after the target depth had been identified. Pipettes had a resistance of 5-8 MΩ and were filled with an internal solution containing (in mM): 115 K-Gluconate, 10 KCl, 10 NaCl, 10 Hepes, 0.1 EGTA, 10 Tris-phosphocreatine, 5 KOH, 13.4 Biocytin, 5 Mg-ATP, 0.3 Tris-GTP. The internal solution had an osmolarity of 300 mOsm and a pH of 7.27 at room temperature. Pipettes are pulled from borosilicate capillaries (OD: 1.0 mm, ID: 0.58 mm; Sutter Instrument Company) using a Model P-2000 puller (Sutter Instrument Company) and inserted into the brain in the coronal plane with a ∼15 degree angle pointing away from the midline. Recordings were made using a Multiclamp 700B amplifier (Molecular Devices). The V_m_ was not corrected for liquid junction potential. Capacitance neutralization was set prior to establishing the GΩ seal. Access resistance was estimated online by fitting the voltage response to hyperpolarizing current steps (see below).

### Signal acquisition

All electrophysiological signal acquisition was performed with custom Labview software (National Instruments) that we developed. Electrophysiological signals were sampled simultaneously at 25 kHz with 24 bit resolution using AC (PXI-4498, internal gain: 30 dB, range: +/- 316 mV) or DC-coupled (PXIe-4492, internal gain: 0 dB, range: +/- 10 V) analog-to-digital data acquisition cards (National instruments) with built-in anti-aliasing filters for extracellular and intracellular/juxtacellular recordings, respectively.

### Histology and imaging

To identify the recorded neurons, histology and imaging were performed, as previously described (Horikawa and Armstrong, 1988; Hulse et al., 2017, 2016). Following the experiment, mice were deeply anesthetized with 5% isoflurane, decapitated, and the brain extracted to 4% PFA. Brains were fixed at 4° C in 4% paraformaldehyde overnight and transferred to 0.01 M (300 mOsm) phosphate buffered saline (PBS) the next day. Up to one week later, brains were sectioned coronally (100 μm) on a vibrating microtome (Leica), permeabilized with 1% Triton X-100 (v/v) in PBS for 1-2 h, and incubated overnight at room temperature in PBS containing avidin-fluorescein (1:200, Vector Laboratories), 5% (v/v) normal horse serum (NHS), and 0.1% Triton X-100. Sections were rinsed in PBS between each step. The next day, sections containing biocytin stained neurons were identified on an inverted epifluorescent microscope (Olympius IX51) for further immunohistochemical processing. Sections underwent immunohistochemical staining against calbindin (CB) and parvalbumin (PV) to aid locating the recorded neurons in the hippocampus. Sections containing biocytin-stained neurons were first incubated in blocking solution containing 5% NHS, 0.25% Triton X-100, and 0.02% (wt/v) sodium azide in PBS. Next, slices were incubated in PBS containing primary antibodies against CB (Rabbit anti-Calbindin D-28k, 1:2000, Swant) and PV (Goat anti-parvalbumin, 1:2000, Swant) overnight. After thorough rinsing in PBS, slices were incubated in PBS containing secondary antibodies CF543 donkey anti-rabbit (1:500, Biotium) and CF633 donkey anti-goat (1:500, Biotium). Processed slices were rinsed and mounted in antifading mounting medium (EverBrite, Biotium). Stained slices were imaged on an inverted confocal laser-scanning microscope (LSM 710 & LSM 880, Zeiss).

## Data Analysis

### V_m_decomposition

Spikes were detected by identifying local maxima in the broadband membrane potential with peak prominence of at least 15 mV and width of at most 10 ms. The subthreshold membrane potential (V_m_) was computed by interpolating the membrane potential over periods with an action potential starting from 3 ms before to 5 ms after the spike peak. The subthreshold V_m_ signal was then low-pass filtered (Parks-McClellan optimal equiripple FIR filter, 250-350 Hz transition band) and downsampled to 2083 Hz. The V_m_ signal was decomposed into fast, slow, and drift components as follows. First, the slow V_m_ component (V_m,slow_) was obtained by median filtering V_m_ with a 1-second window. Next, the fast V_m_ component (V_m,fast_) was obtained as the residual V_m_ after subtraction of V_m,slow_. A drift component (V_m,drift_) was obtained by smoothing V_m,slow_ with a 60-second boxcar kernel. Finally, V_m,slow_ was detrended by subtracting V_m,drift_, which included the resting membrane potential as well as any V_m_ changes on the timescale of minutes (Figure 2 - figure supplement 3). By construction V_m_ = V_m,fast_ + V_m,slow_ + V_m,drift_ and the components contain different frequency bands of the subthreshold membrane potential.

### Ripple detection

Ripples were detected as transient increases in power in the ripple frequency band of the LFP from the probe site located in the CA1 pyramidal cell layer. Ripple power was estimated by band-pass filtering the LFP trace (80-250 Hz), smoothing its square with a Gaussian kernel (10 ms), and taking the square root. Candidate ripple events were identified as segments for which the ripple power was more than 3 s.d. above the mean. Segments that were less than 55 ms apart were merged, and after the merging step segments of length less than 20 ms were rejected as artifacts. A reference recording site away from the CA1 cell layer that does not exhibit ripples was identified and the same procedure was applied for ripple detection on this reference channel. Events detected in both the CA1 and the reference LFP trace were rejected as artifacts.

### Current source density (CSD) estimation

Local field potentials (LFPs) were recorded from a 32-site silicon probe with 100 µm site spacing positioned so that sites spanned all of neocortex, area CA1, the dentate gyrus, and parts of the thalamus. LFPs were bandpass filtered between 1 Hz and 1 KHz and downsampled to 2083 Hz. Channels with recording artifacts were excluded from the CSD analysis. Laminar current source densities were estimated with ∼17 µm resolution using a robust version of the one-dimensional inverse CSD spline method (Pettersen et al., 2006). In particular, the forward matrix (relating CSDs to LFPs) was computed as usual, but the inverse matrix (relating LFPs to CSDs) was computed using ridge regression with a regularization parameter set by a cross-validation procedure. The spatial smoothness of the CSD estimate was automatically controlled by the regularization parameter and therefore no further spatial smoothing was applied. The anatomical laminae in CA1 and DG were determined using a combination of histological reconstruction of the probe track, electrophysiological markers (ripples, sharp waves, dentate spikes), and the CSD covariance structure. DG CSD magnitude was computed by averaging the rectified CSD signals from the vertical extent (∼200 µm) of the suprapyramidal molecular layer of DG. Before the DG CSD magnitude was converted to a z-score, the signal was smoothed with a 1-second median filter.

### Transfer model estimation

Linear transfer models were estimated after taking the z-scored DG CSD magnitude as model input and the slow V_m_ component of the subthreshold membrane potential of a given cell as output. The finite impulse response (FIR) was estimated from the input-output data using a regularized nonparametric procedure (impulseest in Matlab System Identification Toolbox with tuned and correlated ‘TC’ kernel used for regularization) (Chen et al., 2012). The input-output data were first downsampled to ∼20 Hz and the order of the FIR was set to 75, corresponding to 3.6 second duration. The procedure automatically estimated FIR values at negative delays (up to -0.86 sec) and non-zero filter values at negative delays indicated that the slow V_m_ component led the DG CSD magnitude. Once the impulse responses were estimated the corresponding step responses were simulated by feeding the model with a step input. Low order ARX models were also estimated in a similar manner and led to qualitatively similar results as the FIR models (data not shown).

### UP and DOWN state segmentation

UP and DOWN states (UDS) were identified from the z-scored DG molecular layer CSD magnitude using a hidden Markov model (HMM) (McFarland et al., 2011). First, a binary Gaussian mixture was fit to the distribution of DG CSD values (Figure 2C). Next, the mixture components were used to initialize the emission probability distributions of a two state (UP and DOWN) HMM and then the state transition and emission probabilities were estimated from the DG CSD time series data. Finally, the most likely sequence of states given the observed DG CSD time series, downsampled to 4 Hz, were computed with the Viterbi algorithm. The resulting Viterbi path was used to assign a UDS phase to each time point based on its position relative to the nearest UDS transition times. The method was unsupervised and did not require user tuning of any model parameters.

### Adapting recurrent neural population model

When uncoupled, the EC and CA3 population dynamics follow the Wilson-Cowan type r-a model studied in detail in (Levenstein et al., 2019). Briefly, the mean firing rate of the EC population (r_ec_) evolves under activity-driven adaptation (a_ec_) according to the equations:

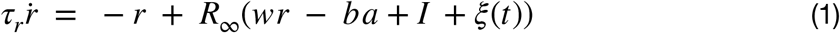

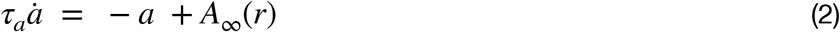

with sigmoid activation functions *R*_∞_(*x*) and *A*_∞_(*x*) given by the logistic curve

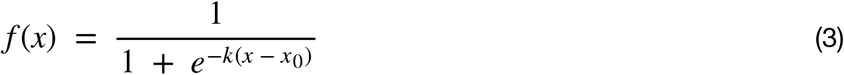

where *k* = 1, *x*_*o*_ = 5 for *R*_∞_(*x*) and *k* = 15, *x*_*o*_ = 0.5 for *A*_∞_(*x*). The time constants *τ*_*r*_ = 1 and *τ*_*a*_ = 25 are dimensionless and set the arbitrary units (AU) of the time axis. For the EC population the strength of recurrent excitation (*w*), adaptation weight (*b*), and the tonic drive (*I*) are all constant parameters with the following values: *w* = 6.8, *b* = 1, I = 2.1. The model is excited by stochastic fluctuations *ξ*(*t*) given by an Ornstein–Uhlenbeck process

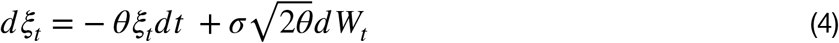

with parameters *θ* = 0.05 and *σ* = 0.1, corresponding to a time constant of 20 and steady-state standard deviation of. The mean firing rate of the CA3 population evolves under the same equations (1-4), but the drive is now a linear function of EC activity

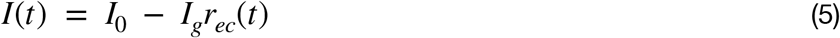

With *I*_0_ = 2.5 and *I*_*g*_ = 0.6, so that elevated EC activity generates a net inhibitory drive to CA3. The strength of recurrent excitation in CA3 is also a linear function of the EC adaptation parameter

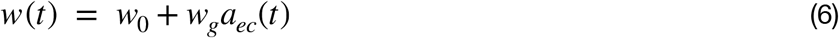

with *w*_0_ = 3.5 and *w*_*g*_ = 2.5, so that the EC UP state leads to an increase in the strength of CA3 recurrent excitation. The stochastic fluctuations exciting the CA3 model have and are unrelated to those in EC.

### Measuring and setting access resistance

Access resistance was estimated online using custom-written software in Labview that communicated with the software (Commander, Molecular Devices) controlling the MultiClamp 700B amplifier through an application programming interface (API). To estimate the access resistance, the bridge balance was temporarily turned off. Then, two -100 pA current pulses (250 ms duration, 250 ms inter-pulse interval) were delivered, the first 50 ms of the hyperpolarizing voltage responses was fit using a simple model, and if the r^2^ fit exceeds 0.99, the bridge balance was set to its new value, otherwise it was returned to the previous value. This procedure was performed once every minute during whole-cell recordings. In addition, all recording parameters in the Commander software were acquired once every second using the API, time stamped to electrophysiological signals, and saved for offline review. The pipette’s voltage response to hyperpolarizing current steps was fit online using a simple double exponential model (Anderson et al., 2000). The computational simplicity of this model sped online fitting. For offline estimates, we used a biophysically-inspired, single-compartment model (Sa et al., 2001). The results obtained from the two models were nearly identical under our recording conditions.

## AUTHOR CONTRIBUTIONS

AGS, EVL, BKH, KK designed the experiments. KK, BKH performed the experiments. EVL led the analysis in collaboration with KK, AGS, BKH. EVL, AGS, KK wrote the paper with input from BKH.

## ACKNOWLEDGEMENTS

We thank Lee-Peng Mok for help with histological processing and immunohistochemistry, Maria Papadopoulou and Stijn Cassenear for help with imaging, insightful discussions and feedback on the manuscript, and Kevin Shan for help with the data processing pipeline and insightful discussion. Confocal imaging was performed at the Caltech Biological Imaging Facility. This work was supported by a Vannevar Bush Faculty Fellowship, the Mathers Foundation, the McKnight Foundation, and NIH grant RO1MH113016.

## Supplementary Figures

**Figure 2 - figure supplement 1.**
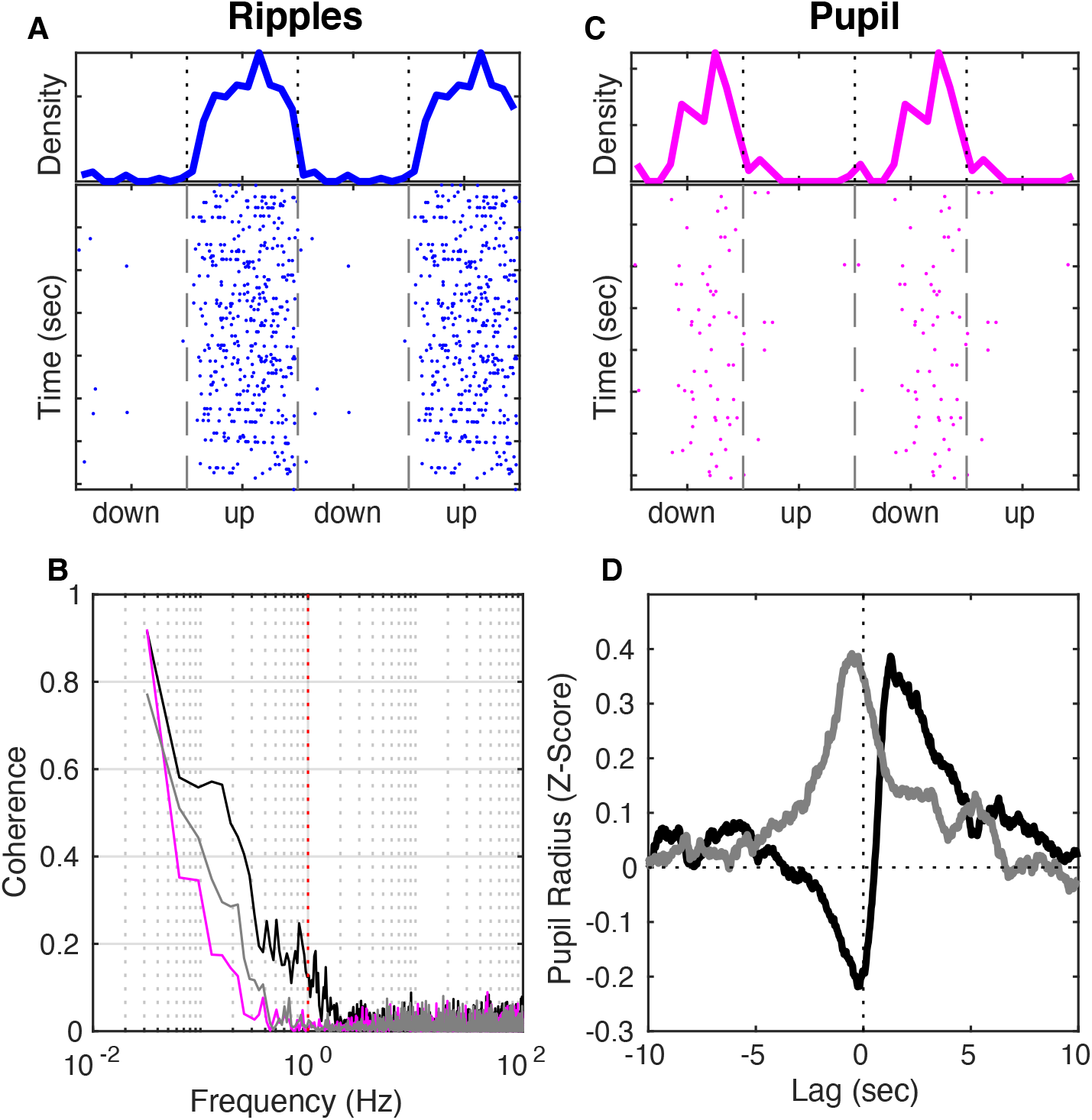
UP and DOWN State Modulation of Ripple and Blink Occurrence. **(A)** (Top) Probability density of ripple occurrence as a function of UDS phase for the recording in Fig. 2. (Bottom) Dots mark the time and phase of each detected ripple. Notice that almost all ripples occur in the UP state. **(B)** Coherence of subthreshold V_m_ and DG CSD magnitude (black) or pupil diameter (magenta) for the recording in Fig. 2. Coherence of DG CSD and pupil diameter in gray. Notice that signals are coherent below 1 Hz. **(C)** Probability density of eye blinks for the same recording session as A. Notice that almost all eye blinks in this session occurred in the DOWN state. **(D)** Pupil radius triggered on transitions to DOWN state (black) and UP state (gray). Notice that the pupil starts dilating at the onset of the DOWN state and is in the process of constricting at the onset of the UP state.

**Figure 2 - figure supplement 2.**
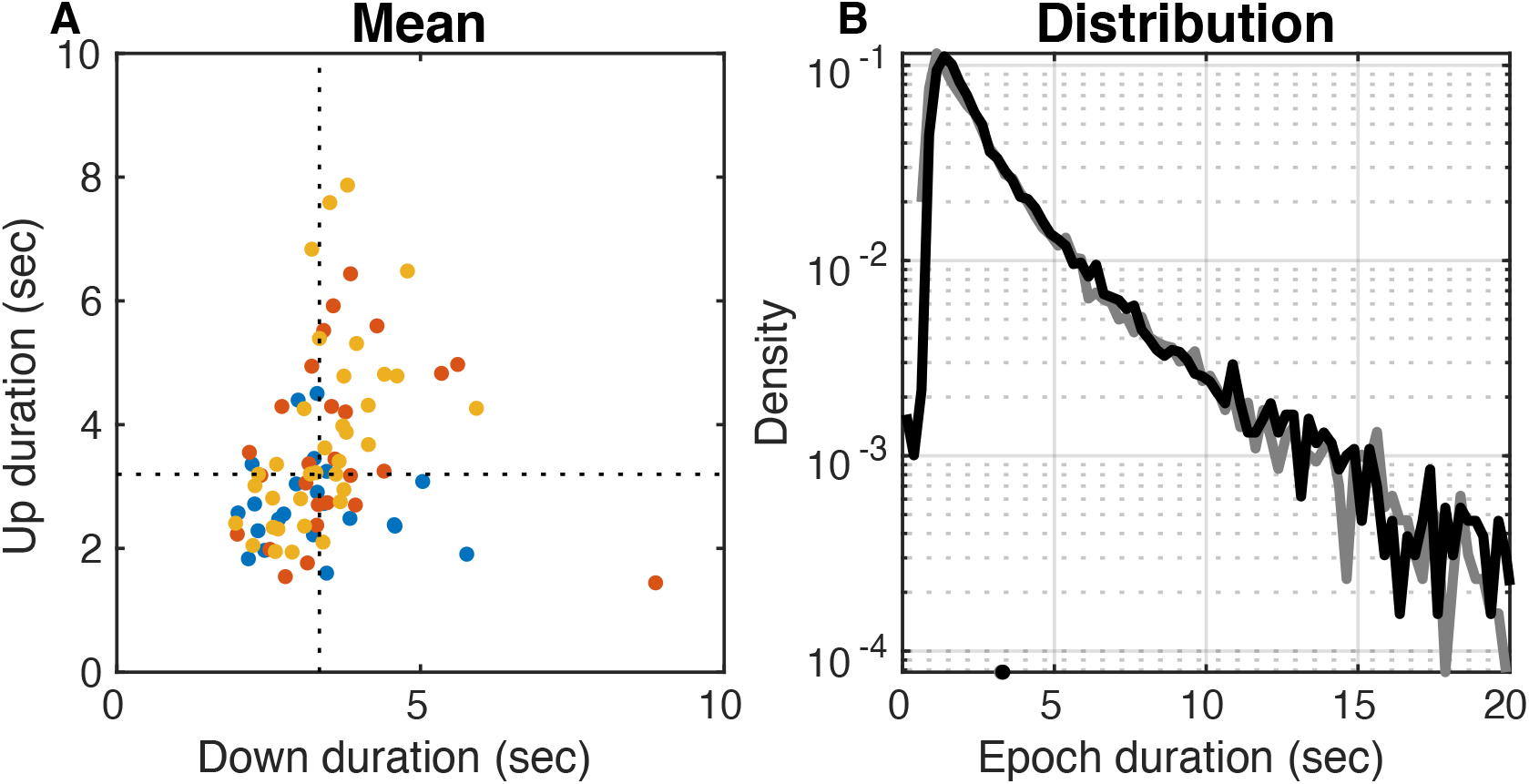
UP and DOWN Epoch Durations. **(A)** Mean DOWN epoch duration plotted against mean UP epoch duration for each dataset color-coded by subfield of whole-cell target. Population averages marked by the interrupted lines (UP = 3.20 sec, DOWN = 3.34 sec). **(B)** Distributions of DOWN (black) and UP (gray) epoch durations from all datasets. Epochs are detected with 250 ms resolution, so the curves at durations shorter than that are affected. Distributions are nearly exponential with similar means (UP = 3.14 sec, DOWN = 3.28 sec) and standard deviations (UP = 3.66 sec, DOWN = 3.56 sec).

**Figure 2 - figure supplement 3.**
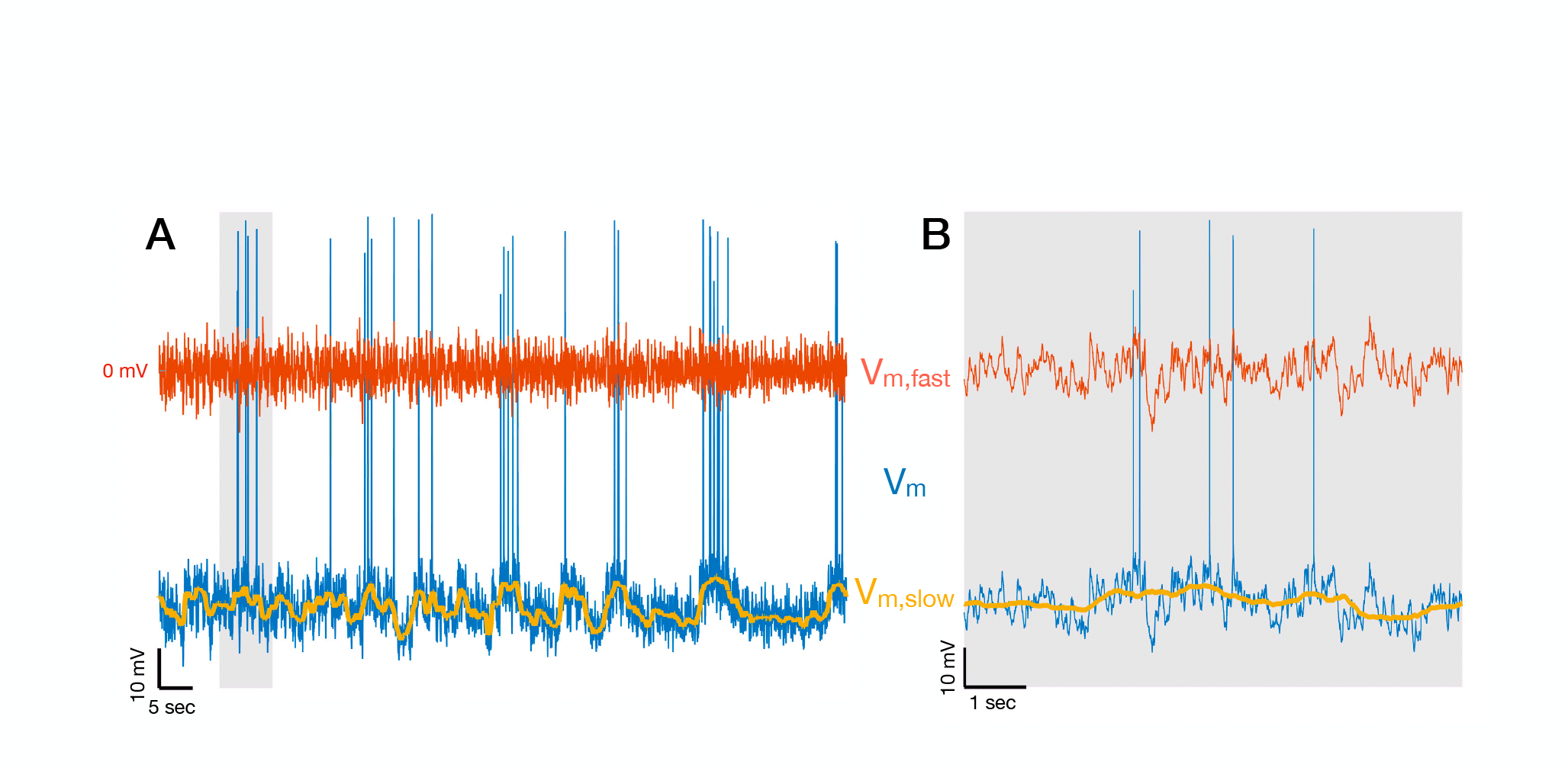
Decomposing Membrane Potential Traces into Slow and Fast Components. **(A)** Membrane potential V_m_ of a CA3 pyramidal cell (blue) decomposed into slow (V_m,slow_ + V_m,drift_ in yellow) and fast (V_m,fast_ in red) components. After spikes are removed from the membrane potential, V_m,slow_ + V_m,drift_ is obtained by filtering V_m_ with a median filter with 1 sec window, and V_m,fast_ = V_m_ - (V_m,slow_ + V_m,drift_). The 1 Hz cutoff frequency for the decomposition is dictated by the fact that V_m_ is coherent with brain state indicators up to 1 Hz (figure 2 - figure supplement 1B). V_m,drift_ contains the resting membrane potential and V_m_ changes on the timescale of minutes, so it only adds a DC offset in the panels above. The short segment indicated by the gray vertical bar is shown in **(B)**.

**Figure 3 - figure supplement 1.**
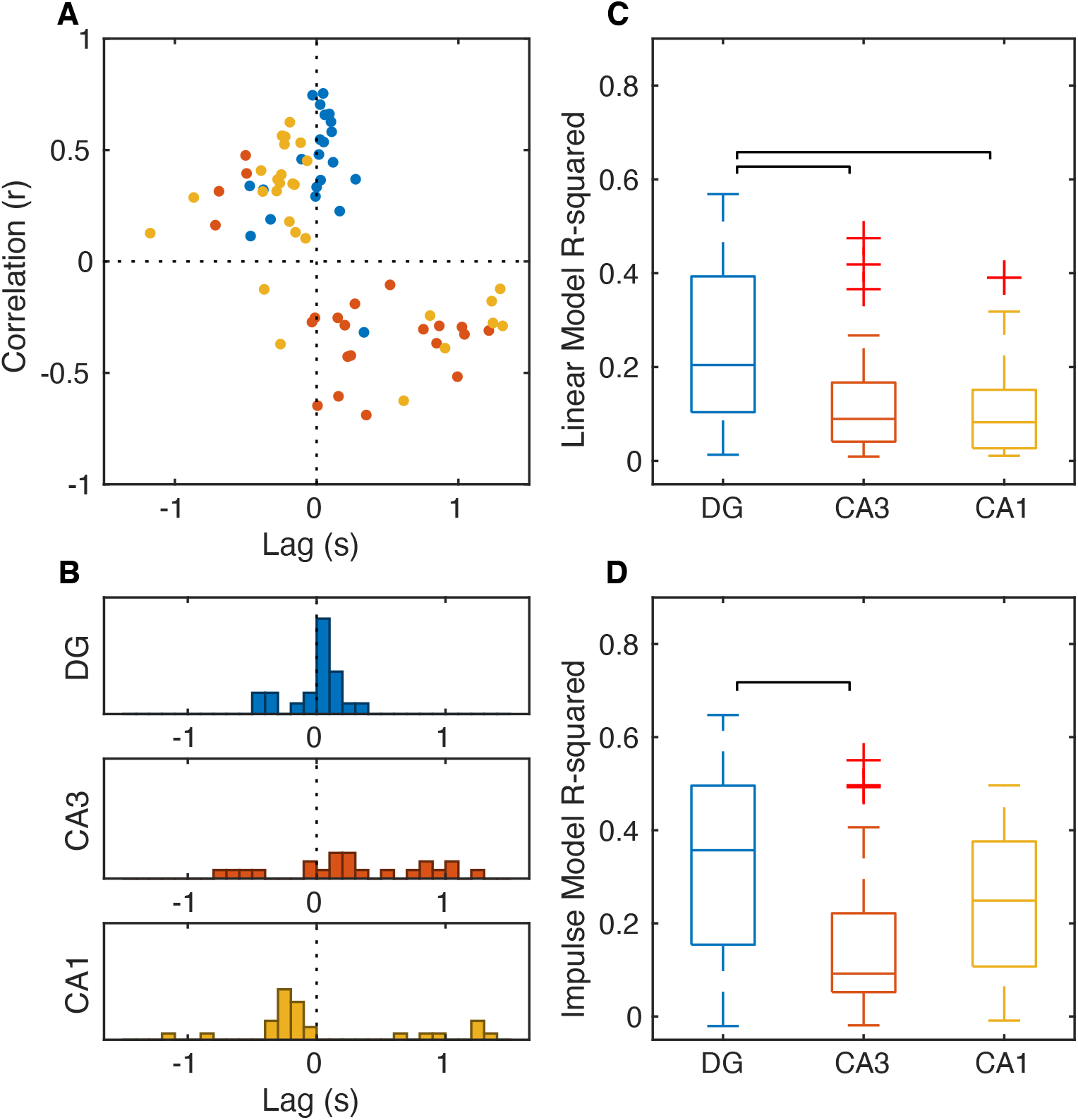
Strength and Direction of the Correlation between V_m_ Slow Component and DG CSD Magnitude. **(A)** Lag and amplitude at the absolute peak of the cross-covariance between V_m_ slow and DG CSD. Each dot represents a cell and is color-coded by subfield. Notice that most DG and CA1 cells have a positive peak correlation, while CA3 is negative. Notice that the peak correlation for many CA1 cells occurs at a negative lag (V_m_ slow leading DG CSD). **(B)** Distribution of peak lags for each subfield. Median lags for cells with significant correlation (|r| > 0.3) for each subfield were: DG (37 ms), CA3 (296 ms), CA1 (−227 ms). **(C)** Fraction of V_m_ slow component variance accounted for by its linear correlation to DG CSD. The medians are DG (20%), CA3 (9%), CA1 (8%). **(D)** Fraction of V_m_ slow component variance that can be explained by an impulse response transfer model with DG CSD as input. The medians are DG (36%), CA3 (9%), CA1 (25%).

**Figure 4 - figure supplement 1.**
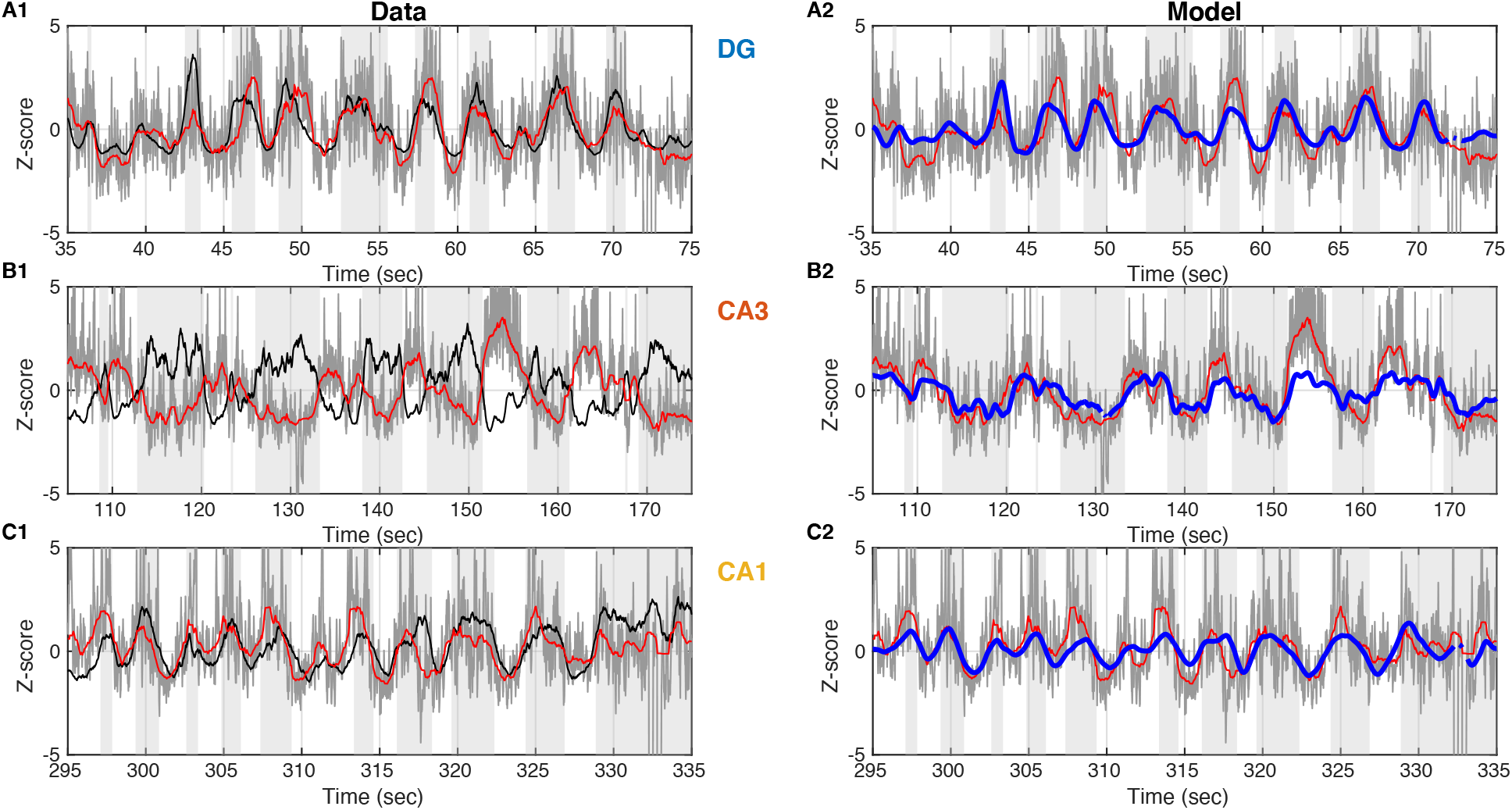
Linear Prediction of Slow V_m_ Component from DG CSD Magnitude. **(A1)** Data from an example DG granule cell. V_m_ (with spikes removed) is shown in gray and its slow component in red. DG CSD is plotted in black and the UP states are marked by the light gray stripes in the background. All traces are converted to z-scores in order to be compared. **(A2)** Same as A1, but the DG CSD trace is omitted and replaced by a linear model prediction of V_m_ slow (blue) from DG CSD. **(B)** Same as A, but for an example CA3 pyramidal cell. **(C)** Same as A and B, but for an example CA1 pyramidal cell.

**Figure 5 - figure supplement 1.**
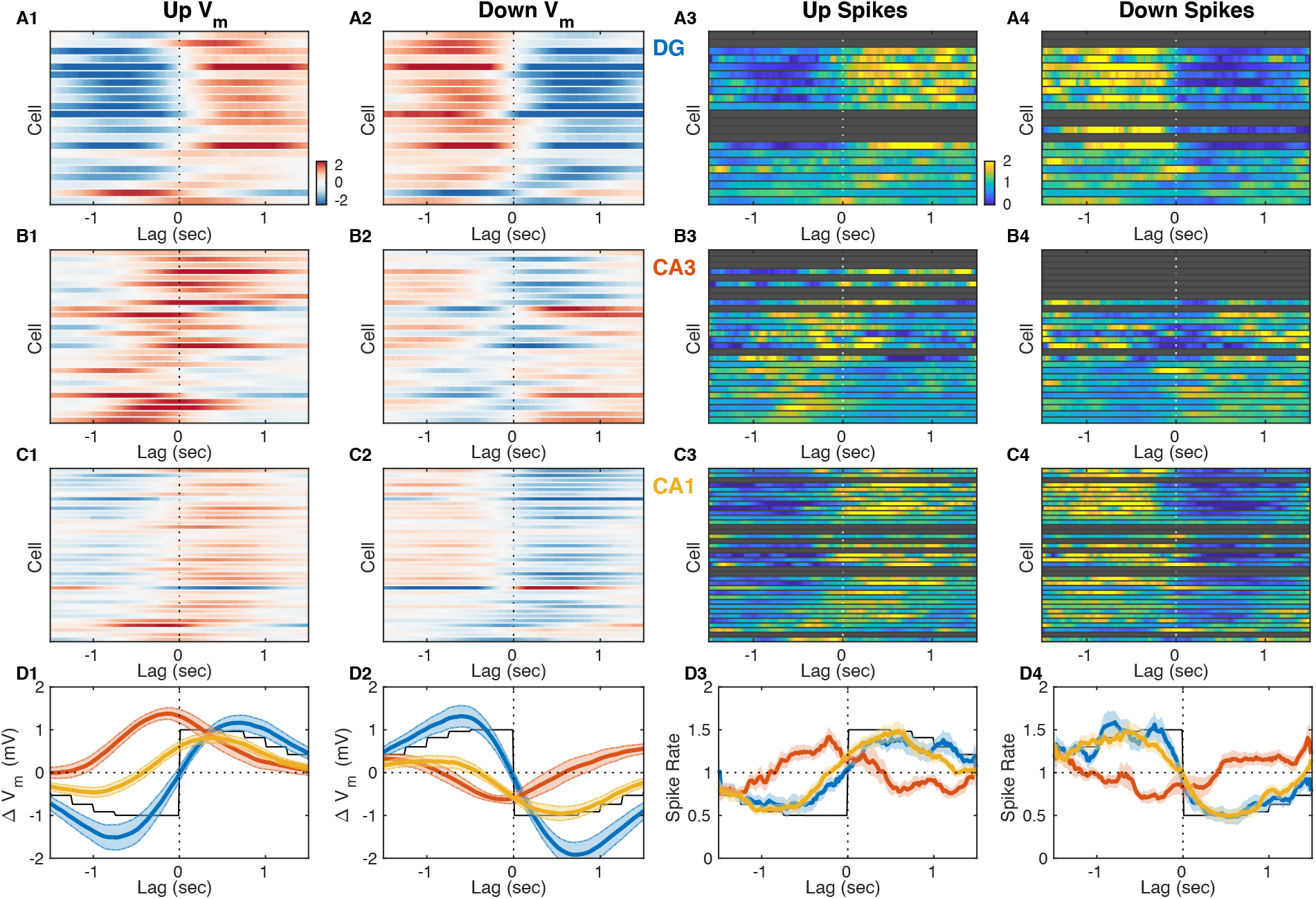
V_m_ and Spiking Responses to UDS Transitions Reveal Ordering in Subfield Activation. **(A1)** Mean slow V_m_ component (V_m,slow_) triggered on DOWN→UP transitions for each DG granule cell displayed as a row in the pseudocolor image. Notice that the V_m_ of most DG granule cells shifts from hyperpolarized (blue) to depolarized (red) at the UP transition (interrupted vertical line). Color limits in all V_m_ panels are ±2.5 mV. **(A2)** Same as A1, but triggered on UP→DOWN transitions. Notice that in DG the V_m_ responses to DOWN state transitions are mostly mirror symmetric to the responses to UP transitions. **(A3)** Spiking responses to DOWN→UP state transitions for all DG granule cells. The perievent time histogram (PETH) for each neuron is normalized to a baseline rate of 1 Hz, so the colors represent the relative modulation of each cell’s baseline firing rate. Notice that firing in the DG is depressed before (blue) and elevated following (yellow) the UP state transition. Grayed out rows correspond to cells with insufficient firing to compute a meaningful PETH. **(A4)** Same as A3, but triggered on UP→DOWN transitions. Notice the sharp decrease of firing right at the onset of the DOWN state. **(B)** Same as A, but for CA3 pyramidal neurons. Notice that many CA3 cells are already depolarized and have increased firing before the onset of the UP transition. **(C)** Same as A and B, but for CA1 pyramidal neurons. **(D)** Area-specific population average V_m_ responses to UP state transitions (D1), DOWN state transitions (D2), and spiking responses to UP transitions (D3) and DOWN transitions (D4). Bands around the mean curves show the standard error of the mean. Black traces show the average state at each lag with the DOWN and UP states represented by -1 and 1 in D1-2, 0.5 and 1.5 in D3-4.

**Figure 5 - figure supplement 2.**
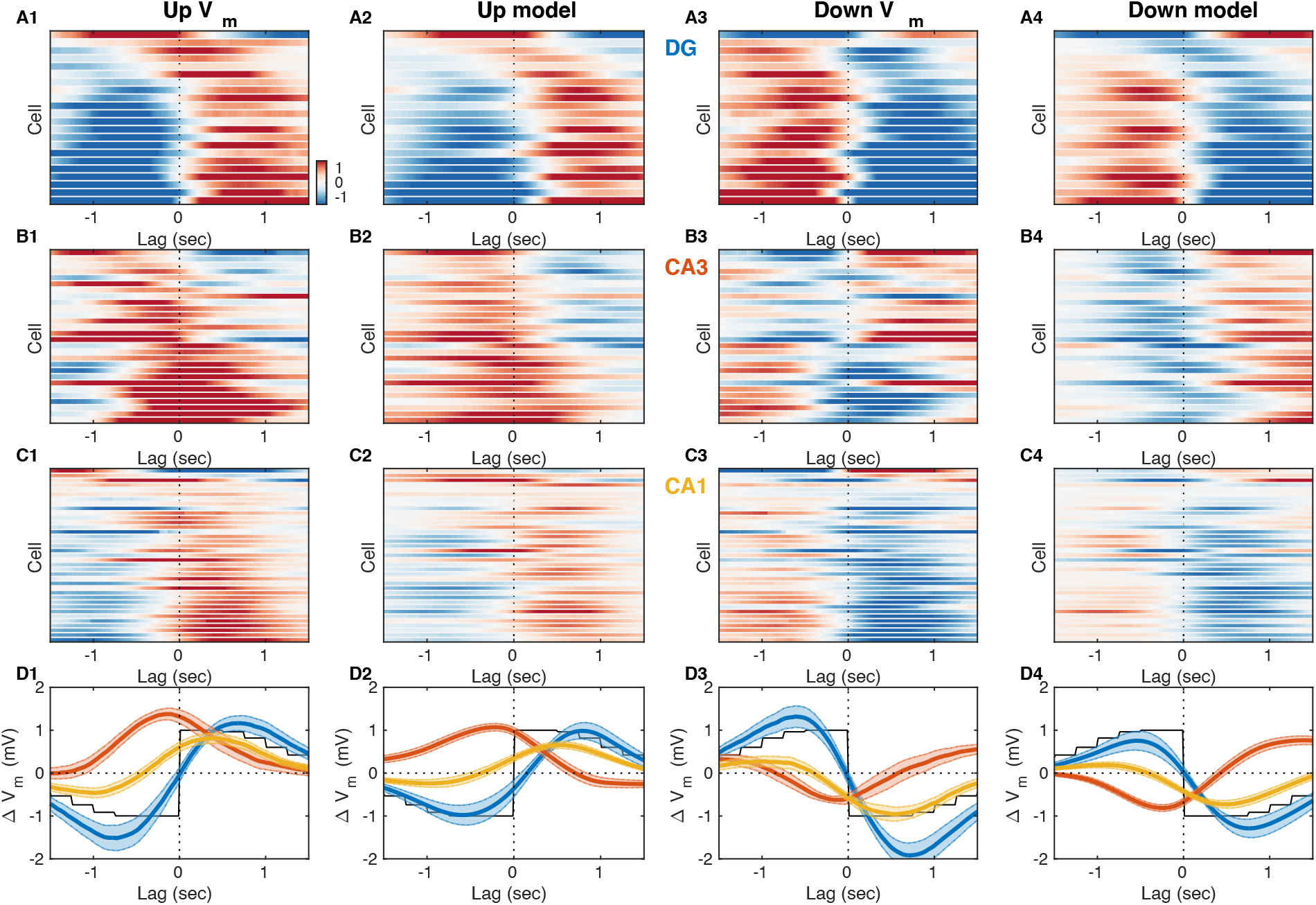
V_m_ and Transfer Model Responses at UP and DOWN State Transitions. **(A1-D1)** Observed V_m_ responses at DOWN→UP transitions. Data are replotted from Fig. 4, but reordered according to the UP V_m_ responses. All other panels follow this same ordering. **(A2-D2)** Transfer model responses at DOWN→UP transitions simulated from the observed DG CSD magnitude. **(A3-D3)** Observed Vm responses at UP→DOWN transitions (replotted from Fig. 4). **(A4-D4)** Transfer model responses at UP→DOWN transitions simulated from the observed DG CSD magnitude.

**Figure 5 - figure supplement 3.**
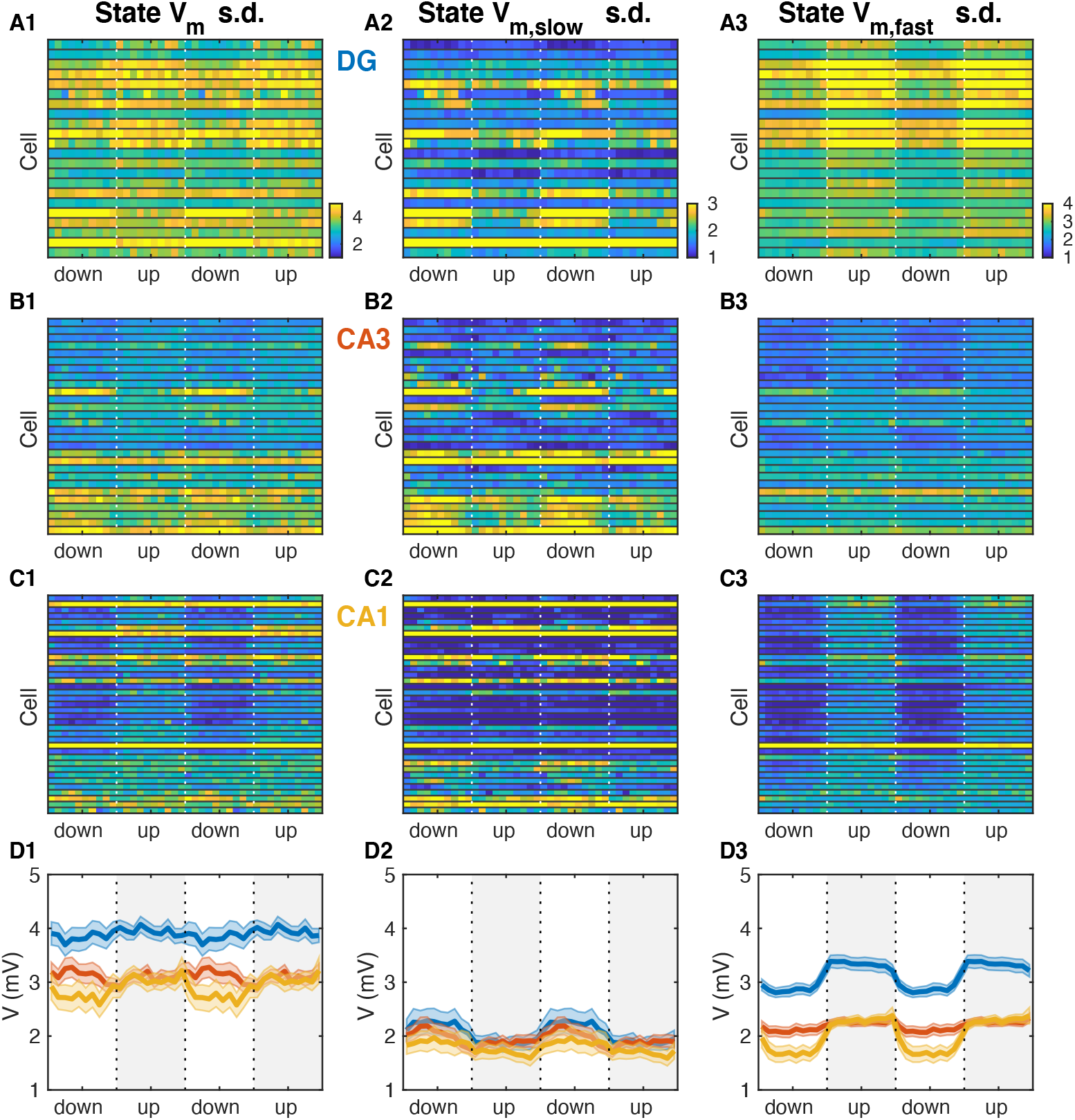
UDS Modulation of V_m_ fluctuations. **(A1)** V_m_ standard deviation (s.d.) as a function of UDS phase for DG granule cells. Vertical interrupted lines mark state transitions and in all panels data are replotted over two UDS cycles. **(A2)** Slow V_m_ component s.d. as a function of UDS phase for DG granule cells. **(A3)** Fast V_m_ component s.d. as a function of UDS phase for DG granule cells. **(B)** Same as A, but for CA3 pyramidal neurons. **(C)** Same as A and B, but for CA1 pyramidal neurons. **(D)** Area-specific population averages color-coded by brain area. (D1) Population average V_m_ s.d. (D2) Population average slow V_m_component s.d. (D3) Population average fast V_m_ component s.d.

## Notes

### Competing Interest Statement

The authors have declared no competing interest.

## References

Acsády L, Kamondi A, Sík A, Freund T, Buzsáki G. 1998. GABAergic cells are the major postsynaptic targets of mossy fibers in the rat hippocampus. J Neurosci 18:3386–3403.

Amaral DG, Witter MP. 1989. The three-dimensional organization of the hippocampal formation: a review of anatomical data. Neuroscience 31:571–591.

Anderson JS, Carandini M, Ferster D. 2000. Orientation tuning of input conductance, excitation, and inhibition in cat primary visual cortex. J Neurophysiol 84:909–926.

Basu J, Zaremba JD, Cheung SK, Hitti FL, Zemelman BV, Losonczy A, Siegelbaum SA. 2016. Gating of hippocampal activity, plasticity, and memory by entorhinal cortex long-range inhibition. Science 351:aaa5694.

Battaglia FP, Sutherland GR, McNaughton BL. 2004. Hippocampal sharp wave bursts coincide with neocortical “up-state” transitions. Learn Mem 11:697–704.

Bazelot M, Teleńczuk MT, Miles R. 2016. Single CA3 pyramidal cells trigger sharp waves in vitro by exciting interneurones. J Physiol 594:2565–2577.

Buzsáki G. 2015. Hippocampal sharp wave-ripple: A cognitive biomarker for episodic memory and planning. Hippocampus 25:1073–1188.

Buzsáki G. 1986. Hippocampal sharp waves: Their origin and significance. Brain Research. doi:10.1016/0006-8993(86)91483-6

Chen T, Ohlsson H, Ljung L. 2012. On the estimation of transfer functions, regularizations and Gaussian processes—Revisited. Automatica. doi:10.1016/j.automatica.2012.05.026

Cowan RL, Wilson CJ. 1994. Spontaneous firing patterns and axonal projections of single corticostriatal neurons in the rat medial agranular cortex. J Neurophysiol 71:17–32.

Davoudi H, Foster DJ. 2019. Acute silencing of hippocampal CA3 reveals a dominant role in place field responses. Nat Neurosci 22:337–342.

de la Prida LM, Huberfeld G, Cohen I, Miles R. 2006. Threshold behavior in the initiation of hippocampal population bursts. Neuron 49:131–142.

Ego-Stengel V, Wilson MA. 2010. Disruption of ripple-associated hippocampal activity during rest impairs spatial learning in the rat. Hippocampus 20:1–10.

Ellender TJ, Nissen W, Colgin LL, Mann EO, Paulsen O. 2010. Priming of hippocampal population bursts by individual perisomatic-targeting interneurons. J Neurosci 30:5979–5991.

English DF, Peyrache A, Stark E, Roux L, Vallentin D, Long MA, Buzsáki G. 2014. Excitation and inhibition compete to control spiking during hippocampal ripples: intracellular study in behaving mice. J Neurosci 34:16509–16517.

Evangelista R, Cano G, Cooper C, Schmitz D, Maier N, Kempter R. 2020. Generation of Sharp Wave-Ripple Events by Disinhibition. J Neurosci 40:7811–7836.

Foster DJ. 2017. Replay Comes of Age. Annu Rev Neurosci 40:581–602.

Girardeau G, Benchenane K, Wiener SI, Buzsáki G, Zugaro MB. 2009. Selective suppression of hippocampal ripples impairs spatial memory. Nat Neurosci 12:1222–1223.

Hahn TTG, McFarland JM, Berberich S, Sakmann B, Mehta MR. 2012. Spontaneous persistent activity in entorhinal cortex modulates cortico-hippocampal interaction in vivo. Nat Neurosci 15:1531–1538.

Hahn TTG, Sakmann B, Mehta MR. 2007. Differential responses of hippocampal subfields to cortical up-down states. Proc Natl Acad Sci U S A 104:5169–5174.

Hasselmo ME, Schnell E. 1994. Laminar selectivity of the cholinergic suppression of synaptic transmission in rat hippocampal region CA1: computational modeling and brain slice physiology. J Neurosci 14:3898–3914.

Hasselmo ME, Schnell E, Barkai E. 1995. Dynamics of learning and recall at excitatory recurrent synapses and cholinergic modulation in rat hippocampal region CA3. J Neurosci 15:5249–5262.

Horikawa K, Armstrong WE. 1988. A versatile means of intracellular labeling: injection of biocytin and its detection with avidin conjugates. J Neurosci Methods 25:1–11.

Hulse BK, Lubenov EV, Siapas AG. 2017. Brain State Dependence of Hippocampal Subthreshold Activity in Awake Mice. Cell Rep 18:136–147.

Hulse BK, Moreaux LC, Lubenov EV, Siapas AG. 2016. Membrane Potential Dynamics of CA1 Pyramidal Neurons during Hippocampal Ripples in Awake Mice. Neuron 89:800–813.

Hunt DL, Linaro D, Si B, Romani S, Spruston N. 2018. A novel pyramidal cell type promotes sharp-wave synchronization in the hippocampus. Nat Neurosci 21:985–995.

Isomura Y, Sirota A, Ozen S, Montgomery S, Mizuseki K, Henze DA, Buzsáki G. 2006. Integration and segregation of activity in entorhinal-hippocampal subregions by neocortical slow oscillations. Neuron 52:871–882.

Jadhav SP, Kemere C, German PW, Frank LM. 2012. Awake hippocampal sharp-wave ripples support spatial memory. Science 336:1454–1458.

Jarosiewicz B, Skaggs WE. 2004. Level of arousal during the small irregular activity state in the rat hippocampal EEG. J Neurophysiol 91:2649–2657.

Jiang X, Gonzalez-Martinez J, Halgren E. 2019. Coordination of Human Hippocampal Sharpwave Ripples during NREM Sleep with Cortical Theta Bursts, Spindles, Downstates, and Upstates. J Neurosci 39:8744–8761.

Ji D, Wilson MA. 2007. Coordinated memory replay in the visual cortex and hippocampus during sleep. Nat Neurosci 10:100–107.

Katona L, Lapray D, Viney TJ, Oulhaj A, Borhegyi Z, Micklem BR, Klausberger T, Somogyi P. 2014. Sleep and movement differentiates actions of two types of somatostatin-expressing GABAergic interneuron in rat hippocampus. Neuron 82:872–886.

Kay K, Sosa M, Chung JE, Karlsson MP, Larkin MC, Frank LM. 2016. A hippocampal network for spatial coding during immobility and sleep. Nature 531:185–190.

Klausberger T, Magill PJ, Márton LF, Roberts JDB, Cobden PM, Buzsáki G, Somogyi P. 2003. Brain-state- and cell-type-specific firing of hippocampal interneurons in vivo. Nature 421:844–848.

Kudrimoti HS, Barnes CA, McNaughton BL. 1999. Reactivation of hippocampal cell assemblies: effects of behavioral state, experience, and EEG dynamics. J Neurosci 19:4090–4101.

Lee AK, Wilson MA. 2002. Memory of sequential experience in the hippocampus during slow wave sleep. Neuron 36:1183–1194.

Levenstein D, Buzsáki G, Rinzel J. 2019. NREM sleep in the rodent neocortex and hippocampus reflects excitable dynamics. Nat Commun 10:2478.

Logothetis NK, Eschenko O, Murayama Y, Augath M, Steudel T, Evrard HC, Besserve M, Oeltermann A. 2012. Hippocampal-cortical interaction during periods of subcortical silence. Nature 491:547–553.

Malezieux M, Kees AL, Mulle C. 2020. Theta Oscillations Coincide with Sustained Hyperpolarization in CA3 Pyramidal Cells, Underlying Decreased Firing. Cell Rep 32:107868.

Margrie TW, Brecht M, Sakmann B. 2002. In vivo, low-resistance, whole-cell recordings from neurons in the anaesthetized and awake mammalian brain. Pflugers Arch 444:491–498.

Marr D. 1971. Simple memory: a theory for archicortex. Philos Trans R Soc Lond B Biol Sci 262:23–81.

McFarland JM, Hahn TTG, Mehta MR. 2011. Explicit-duration hidden Markov model inference of UP-DOWN states from continuous signals. PLoS One 6:e21606.

Mena-Segovia J, Sims HM, Magill PJ, Bolam JP. 2008. Cholinergic brainstem neurons modulate cortical gamma activity during slow oscillations. J Physiol 586:2947–2960.

Middleton SJ, McHugh TJ. 2020. CA2: A Highly Connected Intrahippocampal Relay. Annu Rev Neurosci 43:55–72.

Miles R, Wong RKS. 1983. Single neurones can initiate synchronized population discharge in the hippocampus. Nature. doi:10.1038/306371a0

Mitzdorf U. 1985. Current source-density method and application in cat cerebral cortex: investigation of evoked potentials and EEG phenomena. Physiol Rev 65:37–100.

Mizunuma M, Norimoto H, Tao K, Egawa T, Hanaoka K, Sakaguchi T, Hioki H, Kaneko T, Yamaguchi S, Nagano T, Matsuki N, Ikegaya Y. 2014. Unbalanced excitability underlies offline reactivation of behaviorally activated neurons. Nat Neurosci 17:503–505.

Mölle M, Yeshenko O, Marshall L, Sara SJ, Born J. 2006. Hippocampal sharp wave-ripples linked to slow oscillations in rat slow-wave sleep. J Neurophysiol 96:62–70.

O’Keefe J. 1976. Place units in the hippocampus of the freely moving rat. Experimental Neurology. doi:10.1016/0014-4886(76)90055-8

Oliva A, Fernández-Ruiz A, Buzsáki G, Berényi A. 2016. Role of Hippocampal CA2 Region in Triggering Sharp-Wave Ripples. Neuron 91:1342–1355.

Pettersen KH, Devor A, Ulbert I, Dale AM, Einevoll GT. 2006. Current-source density estimation based on inversion of electrostatic forward solution: effects of finite extent of neuronal activity and conductivity discontinuities. J Neurosci Methods 154:116–133.

Pinault D. 1996. A novel single-cell staining procedure performed in vivo under electrophysiological control: morpho-functional features of juxtacellularly labeled thalamic cells and other central neurons with biocytin or Neurobiotin. J Neurosci Methods 65:113–136.

Roumis DK, Frank LM. 2015. Hippocampal sharp-wave ripples in waking and sleeping states. Curr Opin Neurobiol 35:6–12.

Sa VR de, de Sa VR, MacKay DJC. 2001. Model fitting as an aid to bridge balancing in neuronal recording. Neurocomputing. doi:10.1016/s0925-2312(01)00525-2

Sasaki T, Piatti VC, Hwaun E, Ahmadi S, Lisman JE, Leutgeb S, Leutgeb JK. 2018. Dentate network activity is necessary for spatial working memory by supporting CA3 sharp-wave ripple generation and prospective firing of CA3 neurons. Nat Neurosci 21:258–269.

Scharfman HE. 2007. The CA3 “backprojection” to the dentate gyrus. Prog Brain Res 163:627–637.

Schlingloff D, Káli S, Freund TF, Hájos N, Gulyás AI. 2014. Mechanisms of sharp wave initiation and ripple generation. J Neurosci 34:11385–11398.

Shein-Idelson M, Ondracek JM, Liaw H-P, Reiter S, Laurent G. 2016. Slow waves, sharp waves, ripples, and REM in sleeping dragons. Science 352:590–595.

Siapas AG, Wilson MA. 1998. Coordinated interactions between hippocampal ripples and cortical spindles during slow-wave sleep. Neuron 21:1123–1128.

Somogyi P, Katona L, Klausberger T, Lasztóczi B, Viney TJ. 2014. Temporal redistribution of inhibition over neuronal subcellular domains underlies state-dependent rhythmic change of excitability in the hippocampus. Philos Trans R Soc Lond B Biol Sci 369:20120518.

Squire LR. 1992. Memory and the hippocampus: a synthesis from findings with rats, monkeys, and humans. Psychol Rev 99:195–231.

Stark E, Roux L, Eichler R, Senzai Y, Royer S, Buzsáki G. 2014. Pyramidal cell-interneuron interactions underlie hippocampal ripple oscillations. Neuron 83:467–480.

Steriade M, McCormick D, Sejnowski T. 1993a. Thalamocortical oscillations in the sleeping and aroused brain. Science. doi:10.1126/science.8235588

Steriade M, Nunez A, Amzica F. 1993b. A novel slow (< 1 Hz) oscillation of neocortical neurons in vivo: depolarizing and hyperpolarizing components. The Journal of Neuroscience. doi:10.1523/jneurosci.13-08-03252.1993

Steriade M, Nunez A, Amzica F. 1993c. Intracellular analysis of relations between the slow (< 1 Hz) neocortical oscillation and other sleep rhythms of the electroencephalogram. The Journal of Neuroscience. doi:10.1523/jneurosci.13-08-03266.1993

Steward O, Cotman C, Lynch G. 1976. A quantitative autoradiographic and electrophysiological study of the reinnervation of the dentate gyrus by the contralateral entorhinal cortex following ipsilateral entorhinal lesions. Brain Research. doi:10.1016/0006-8993(76)90665-x

Sullivan D, Csicsvari J, Mizuseki K, Montgomery S, Diba K, Buzsáki G. 2011. Relationships between hippocampal sharp waves, ripples, and fast gamma oscillation: influence of dentate and entorhinal cortical activity. J Neurosci 31:8605–8616.

Szabo GG, D. X, Oijala M, Varga C, Parent JM, Soltesz I. 2017. Extended Interneuronal Network of the Dentate Gyrus. Cell Rep 20:1262–1268.

Tamamaki N, Nojyo Y. 1993. Projection of the entorhinal layer II neurons in the rat as revealed by intracellular pressure-injection of neurobiotin. Hippocampus 3:471–480.

Tang W, Jadhav SP. 2019. Sharp-wave ripples as a signature of hippocampal-prefrontal reactivation for memory during sleep and waking states. Neurobiol Learn Mem 160:11–20.

Traub RD, Miles R. 1991. Neuronal Networks of the Hippocampus. Cambridge University Press.

Tukker JJ, Lasztóczi B, Katona L, Roberts JDB, Pissadaki EK, Dalezios Y, Márton L, Zhang L, Klausberger T, Somogyi P. 2013. Distinct dendritic arborization and in vivo firing patterns of parvalbumin-expressing basket cells in the hippocampal area CA3. J Neurosci 33:6809–6825.

Unal G, Crump MG, Viney TJ, Éltes T, Katona L, Klausberger T, Somogyi P. 2018. Spatio-temporal specialization of GABAergic septo-hippocampal neurons for rhythmic network activity. Brain Struct Funct 223:2409–2432.

Vandecasteele M, Varga V, Berényi A, Papp E, Barthó P, Venance L, Freund TF, Buzsáki G. 2014. Optogenetic activation of septal cholinergic neurons suppresses sharp wave ripples and enhances theta oscillations in the hippocampus. Proc Natl Acad Sci U S A 111:13535–13540.

Vanderwolf CH. 1969. Hippocampal electrical activity and voluntary movement in the rat. Electroencephalogr Clin Neurophysiol 26:407–418.

Viney TJ, Lasztoczi B, Katona L, Crump MG, Tukker JJ, Klausberger T, Somogyi P. 2013. Network state-dependent inhibition of identified hippocampal CA3 axo-axonic cells in vivo. Nat Neurosci 16:1802–1811.

Vogt KE, Regehr WG. 2001. Cholinergic modulation of excitatory synaptic transmission in the CA3 area of the hippocampus. J Neurosci 21:75–83.

Wierzynski CM, Lubenov EV, Gu M, Siapas AG. 2009. State-dependent spike-timing relationships between hippocampal and prefrontal circuits during sleep. Neuron 61:587–596.

Wilson M, McNaughton B. 1994. Reactivation of hippocampal ensemble memories during sleep. Science. doi:10.1126/science.8036517

Yamamoto J, Tonegawa S. 2017. Direct Medial Entorhinal Cortex Input to Hippocampal CA1 Is Crucial for Extended Quiet Awake Replay. Neuron 96:217–227.e4.

Zucca S, Griguoli M, Malézieux M, Grosjean N, Carta M, Mulle C. 2017. Control of Spike Transfer at Hippocampal Mossy Fiber Synapses In Vivo by GABAA and GABAB Receptor-Mediated Inhibition. J Neurosci 37:587–598.

